# CCG-1423-derived compounds reduce global RNA synthesis and inhibit transcriptional responses

**DOI:** 10.1101/2023.09.08.556810

**Authors:** Bina Prajapati, Maria Sokolova, Ekaterina Sidorenko, Mikael Kyriacou, Salla Kyheröinen, Anniina Vihervaara, Maria K. Vartiainen

**Affiliations:** Institute of Biotechnology, HiLIFE, University of Helsinki, Finland; Department of Gene Technology, KTH Royal Institute of Technology, Science for Life Laboratory, Stockholm, Sweden

**Author notes:** equal contribution (Bina Prajapati, Maria Sokolova).

## Abstract

Myocardin-related transcription factors (MRTFs) are coactivators of serum response factor (SRF), and thereby regulate cytoskeletal gene expression in response to actin dynamics. MRTFs have also been implicated in heat shock protein (*hsp*) transcription in fly ovaries, but the mechanisms remain unclear. Here we demonstrate that in mammalian cells, MRTFs are dispensable for *hsp* gene induction. However, the widely used small molecule inhibitors of MRTF/SRF transcription pathway, derived from CCG-1423, efficiently inhibit *hsp* gene transcription in both fly and mammalian cells also in absence of MRTFs. Quantifying RNA synthesis and RNA polymerase distribution demonstrates that CCG-1423-derived compounds have a genome-wide effect on transcription. Indeed, tracking nascent transcription at nucleotide resolution reveals that CCG-1423-derived compounds reduce RNA polymerase II elongation, and severely dampen the transcriptional response to heat shock. The effects of CCG-1423-derived compounds therefore extend beyond the MRTF/SRF pathway into nascent transcription, opening novel opportunities for their use in transcription research.

## Introduction

Activation of different transcriptional programs is essential for cellular responses to various internal and external cues, thereby underlying several essential biological processes. Serum response factor (SRF) is an extensively studied transcription factor that regulates the expression of immediate-early, cytoskeletal and muscle-specific genes. As its name implies, SRF-mediated transcription is strongly activated by different components of serum that culminate on two transcription cofactor families, ternary complex factors (TCFs) and myocardin-related transcription factors (MRTFs). These cofactor families compete for binding to SRF, are regulated by distinct signalling pathways and control different cellular processes (Posern and Treisman, 2006). The MRTF family is essential for the regulation of contractile and cytoskeletal target genes via SRF in most tissues and consists of ubiquitously expressed MRTF-A and MRTF-B, as well as heart and smooth muscle-specific myocardin. While the overall domain structure is well-conserved between the family members, the RPEL domain of myocardin does not bind actin. In MRTF-A and MRTF-B, the RPEL domain binds actin, and regulates the subcellular localization and activity of these transcription cofactors in response to actin dynamics. Consequently, myocardin is constitutively nuclear and active, whereas MRTF-A and MRTF-B shuttle in and out of the nucleus in response to changes in the actin-monomer pool (Guettler et al., 2008; Vartiainen et al., 2007).

Actin is one of the transcriptional targets of the MRTF-SRF pathway, creating a feedback loop, where actin dynamics regulate the expression of its components. This feature seems to be evolutionarily conserved. During border cell migration, cell stretching and activation of Diaphanous formin leads to actin polymerization, which induces nuclear accumulation and activation of *Drosophila* MRTF (MAL-D). Here MAL-D/SRF activity is required for building a robust actin cytoskeleton, and thereby for border cell migration (Somogyi and Rorth, 2004). A combination of chromatin immunoprecipitation (ChIP) analysis for MAL-D chromatin binding sites with gene expression analysis of a MAL-D mutant (mal-d^Δ7^) ovaries revealed only a few genes as “direct” MAL-D targets. One of them was the actin gene *Act5C*. Importantly, re-expression of Act5C in MAL-D mutant cells rescued the border cell migration defect (Salvany et al., 2014), attesting to the biological significance of MRTFs in transcriptional regulation of actin. Curiously, the other MAL-D target genes identified in this study were a subset of heat shock protein (*hsp*) genes (Salvany et al., 2014). Heat shock (HS) response is an evolutionary conserved mechanism that triggers production of molecular chaperones to maintain protein homeostasis upon various forms of stress. HS causes a global reprogramming of transcription (reviewed in (Vihervaara et al., 2018)); thousands of genes are repressed upon HS, but simultaneously, hundreds of genes, including several *hsp* genes encoding for classical chaperones, are rapidly activated. The heat-induced genes overcome the repressive environment by recruiting strong trans-activators, such as Heat shock factor 1. Precise mapping of positions and orientations of transcriptionally engaged Pol II during HS, have suggested that in mouse and human cells, inhibition of pause-release causes global redistribution of Pol II, clearing elongating Pol II from the gene bodies of repressed genes (Mahat et al., 2016a; Vihervaara et al., 2017). This genome-wide repression may be facilitated by recruitment of negative elongation factor (NELF) to gene promoters through biomolecular condensation (Rawat et al., 2021). In addition to pause-release, HS has also been shown to reduce processivity of Pol II, resulting in premature termination of transcription (Cugusi et al., 2022) and read-through transcription (Vilborg et al., 2017). Intriguingly, also some cytoskeletal genes, which are SRF targets, have been shown to be transiently activated in response to HS (Mahat and Lis, 2017; Mahat et al., 2016b). The role of MRTFs in this context has not been studied, but it is tempting to speculate that changes in the actin cytoskeleton known to take place in response to HS (Welch and Suhan, 1985) could regulate MRTF localization and activity here similarly as during serum stimulation (Miralles et al., 2003; Vartiainen et al., 2007). Nevertheless, how this links to the plausible role of MRTFs in regulating *hsp* gene expression (Salvany et al., 2014) remains unclear.

Due to the importance of MRTF/SRF for several actin-dependent cellular processes, and altered MRTF/SRF activity associated with various diseases, such as cancer (Brandt et al., 2009) (Gau et al., 2022; Medjkane et al., 2009), there has been considerable interest in developing small molecule inhibitors targeting specifically this transcriptional pathway. A transcription-based high-throughput assay that utilised a serum response element-controlled luciferase reporter was used to identify CCG-1423 as an inhibitor of Rho-activated transcription (Evelyn et al., 2007). CCG-1423, and its improved variants, including CCG-203971 (Johnson et al., 2014) have been widely used in several different experimental systems from cell lines to animal models, and reported to inhibit MRTF/SRF target gene expression, impair cell motility (Bell et al., 2013; Haak et al., 2017; Minami et al., 2012; Zhang et al., 2022; Zhao et al., 2020) and prevent fibrosis in a preclinical model (Sisson et al., 2015; Yu-Wai-Man et al., 2017). Nevertheless, the primary target(s) for these compounds have remained controversial, and it has, for example, been proposed that the target would be MRTF itself (Hayashi et al., 2014), an actin-binding protein MICAL2 (Lundquist et al., 2014) or most recently an iron-binding transcription factor pirin (Lisabeth et al., 2019). Consequently, the mechanisms by which CCG-1423-derived inhibitors influence MRTF/SRF activity have remained enigmatic.

Here we demonstrate that the CCG-1423-derived compounds robustly inhibit *hsp* gene activation upon HS. However, this effect is not dependent on the presence of MRTF transcription factors, indicating a more general role for CCG-1423-derived compounds on transcription than previously acknowledged. In fact, we demonstrate that CCG-1423-derived compounds have a global effect on engaged RNA polymerase II, and thereby on transcription programs in both fly and mammalian cells. CCG-1423-derived compounds reduce RNA synthesis and prevent transcriptional reprogramming upon stress, opening new avenues for these compounds in transcriptional and pharmacological studies.

## Results

### MAL-D is essential for basal and heat-induced *hsp* transcription in fly ovaries

Previous studies have shown that *Drosophila* MRTF (MAL-D) binds five small *hsp* genes and is required for their expression during fly oogenesis (Salvany et al., 2014). However, since *hsp* genes are expressed in fly ovaries already prior to heat shock (HS), this study did not address the potential role of MAL-D in HS-induced transcriptional responses. To study this, we examined *hsp* gene expression in homozygous and heterozygous MAL-D knockout (mal-d^Δ7^) (Somogyi and Rorth, 2004) ovaries, with or without 20 min HS at 37°C. As expected and reported before (Somogyi and Rorth, 2004), the homozygous flies did not express any MAL-D mRNA that corresponded to the deleted region (Figure S1A), and also showed decreased mRNA expression from the actin gene *Act5C* (Figure S1A). While heterozygous flies displayed a robust transcriptional induction of both *Hsp70Aa* and *Hsp68* genes upon HS, the homozygous MAL-D knockout flies showed a greatly reduced response (Figure 1A). This demonstrates that at least in fly ovaries, MAL-D is required for efficient transcriptional induction of *hsp* genes upon HS.

**Figure 1:**
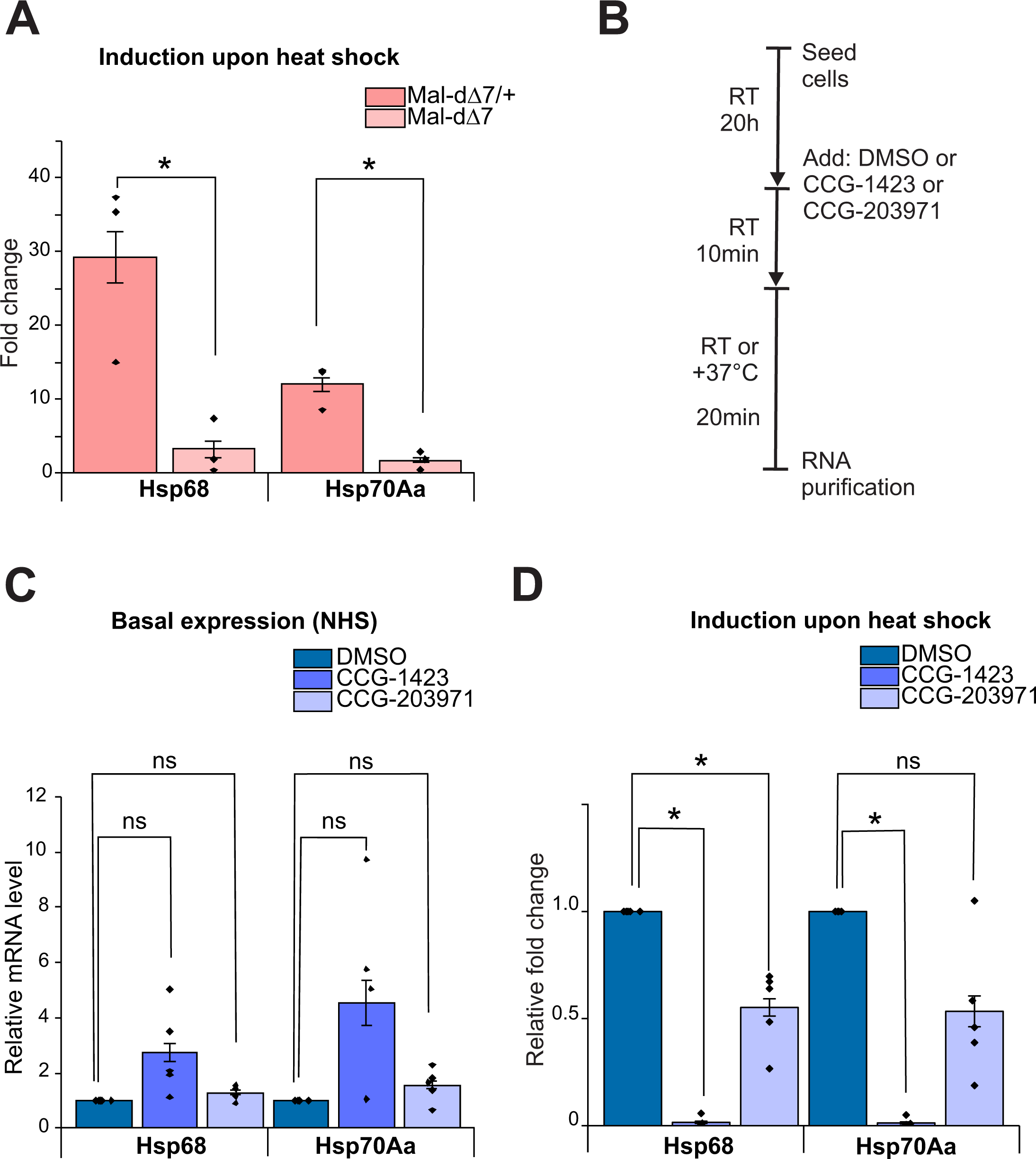
Deletion of MAL-D and CCG-1423-derived inhibitors reduce *hsp* gene induction upon HS. **A.** Deletion of MAL-D reduces transcriptional induction of *Hsp68* and *Hsp70Aa* genes in fly ovaries upon 20 min of heat shock at 37°C. Data is shown as fold changes in *Hsp68* and *Hsp70Aa* mRNA induction in MAL-D-deleted (mal-dΔ7) and heterozygous (mal-dΔ7/+) flies. Data is from three biological replicates, with bar indicating the mean, individual data points shown and error bars are standard error of the mean (s.e.m). Statistical significance with student’s t-test (*; P< 0.05). P-values: Mal-dΔ7/+ vs Mal-dΔ7 for *Hsp68* is 0.0249, Mal-dΔ7/+ vs Mal-dΔ7 for *Hsp70Aa* is 0.0054. **B.** Schematic representation of the experimental design in S2R+ cells for data shown in C and D. **C.** Effect of 30 min treatment with CCG-1423-derived inhibitors on baseline expression (non-heat shock; NH) of *Hsp68* and *Hsp70Aa*. Data is from five biological replicates, normalized to DMSO treated sample and shown as in A. Statistical significance with student’s one-sample t-test (*; P< 0.025 with Bonferroni correction). P-values: CCG-1423 vs DMSO for *Hsp68* is 0.066, CCG-203971 vs DMSO for *Hsp68* is 0.064, CCG-1423 vs DMSO for *Hsp70Aa* is 0.096 and CCG-203971 vs DMSO is 0.108. **D.** CCG-1423 strongly inhibits HS-induced activation of *Hsp68* and *Hsp70Aa* expression. Data is shown as fold change in mRNA induction, from five biological replicates and shown as in A. Statistical significance with student’s one-sample t-test (*; P< 0.025 with Bonferroni correction). P-values: CCG-1423 vs DMSO for *Hsp68* is 8E-8, CCG-203971 vs DMSO for *Hsp68* is 0.005, CCG-1423 vs DMSO for *Hsp70Aa* is 5E-8 and CCG-203971 vs DMSO is 0.032. See also Supplementary Figure 1 for additional data.

### Short exposure to CCG-1423-derived inhibitors decreases *hsp* activation in fly cells

To further explore how MAL-D/MRTF drive *hsp* expression, we used the cultured fly cell line S2R+ with RNAi-mediated knockdown of MAL-D expression. Unfortunately, none of the double stranded RNAs (dsRNAs) targeting distinct MAL-D regions was able to deplete MAL-D mRNA expression below 25% of its basal level (Fig S1B), and did not reproducibly influence HS-induced *Hsp70Aa* transcription compared to control dsRNA targeting GFP (Fig S1C). Lack of antibodies for MAL-D prevented us from measuring protein levels upon RNAi. Hence, we turned to the CCG-1423-derived inhibitors, which have been used in several model systems to inhibit MRTF/SRF-mediated transcription as discussed above, and treated S2R+ cells with CCG-1423 and CCG-203971. When using concentrations and treatment time of 16 h, compatible with those most frequently used in the literature (Evelyn et al., 2007; Hayashi et al., 2014; Sisson et al., 2015), we observed decreased expression of the *Act5C* pre-mRNA (Fig S1D), as would be expected for MRTF/SRF inhibitors. Interestingly, under these conditions, CCG-1423, the original compound, induced the expression of *Hsp70Aa* mRNA 10-fold, and *Hsp68* mRNA to a lesser extent, already in non-HS condition (Fig S1E). Remarkably, CCG-1423 also efficiently prevented transcriptional induction of these genes upon HS (20 min, 37°C; Fig S1F). The second generation derivative CCG-203971 had negligible effects on basal *Hsp70Aa* and *Hsp68* mRNA expression, but inhibited their transcriptional induction upon HS, albeit less than CCG-1423 (Fig S1E,F).

Since 16 h treatment with CCG-compounds caused *hsp* induction in the absence of HS, perhaps due to cellular stress, we next tested shorter incubation times (experimental set up in 1B). Indeed, 30 min treatment with CCG-1423 did not cause significant increase in *hsp* mRNA expression in non-HS cells (Fig 1C), although the trend was similar to the 16 h treatment (Fig S1E). However, already this short incubation with CCG-1423 was sufficient to robustly inhibit HS-triggered induction of *Hsp68* and *Hsp70Aa* genes (Fig 1D), indicating a rapid change in inducibility of transcription. Of note is that 30 min CCG-1423 treatment did not significantly affect *Act5c* pre-mRNA levels (Fig S1G). The effects of CCG-203971 were negligible on both *Act5c* and *hsp* genes (Fig S1G, 1C-D). As a conclusion, CCG-1423 robustly inhibits transcriptional induction of *hsp* genes even with short incubation times, while the second-generation derivative of this compound, CCG-203971, had less of an effect in the cultured fly cells used here.

### MRTFs are not required for *hsp* gene induction in mammalian cells

Next we studied whether MRTFs drive *hsp* gene expression also in mammalian cells. Since MAL-D was shown to bind to *hsp* genes in fly ovaries (Salvany et al., 2014), we first used chromatin immunoprecipitation with deep sequencing (ChIP-seq) in NIH 3T3 mouse fibroblasts to study MRTF chromatin-binding in the presence and absence of HS. However, we did not detect significant enrichment of MRTF near any *hsp* gene promoter in NIH 3T3 cells in either condition (Fig 2A, S2A). Importantly, MRTF clearly interacted with canonical MRTF/SRF target genes, such as *Actb* (Fig S2B), and the overall MRTF chromatin binding pattern was highly similar to published results by us and others (Esnault et al., 2014; Sidorenko et al., 2022). Moreover, RNA polymerase II displayed clear enrichment on *hsp* genes upon HS (Fig 2A, S2A), while MRTF binding to its target genes was reduced upon HS (Fig. S2B), demonstrating both transcriptional activation and repression responses, respectively, to HS, as documented extensively before (Duarte et al., 2016; Himanen et al., 2022; Mahat et al., 2016b; Vihervaara et al., 2017; Vihervaara et al., 2021). Collectively, these results demonstrate that both MRTF ChIP-seq and HS worked as expected, although we failed to detect binding of MRTF to *hsp* gene promoters in mammalian cells.

**Figure 2:**
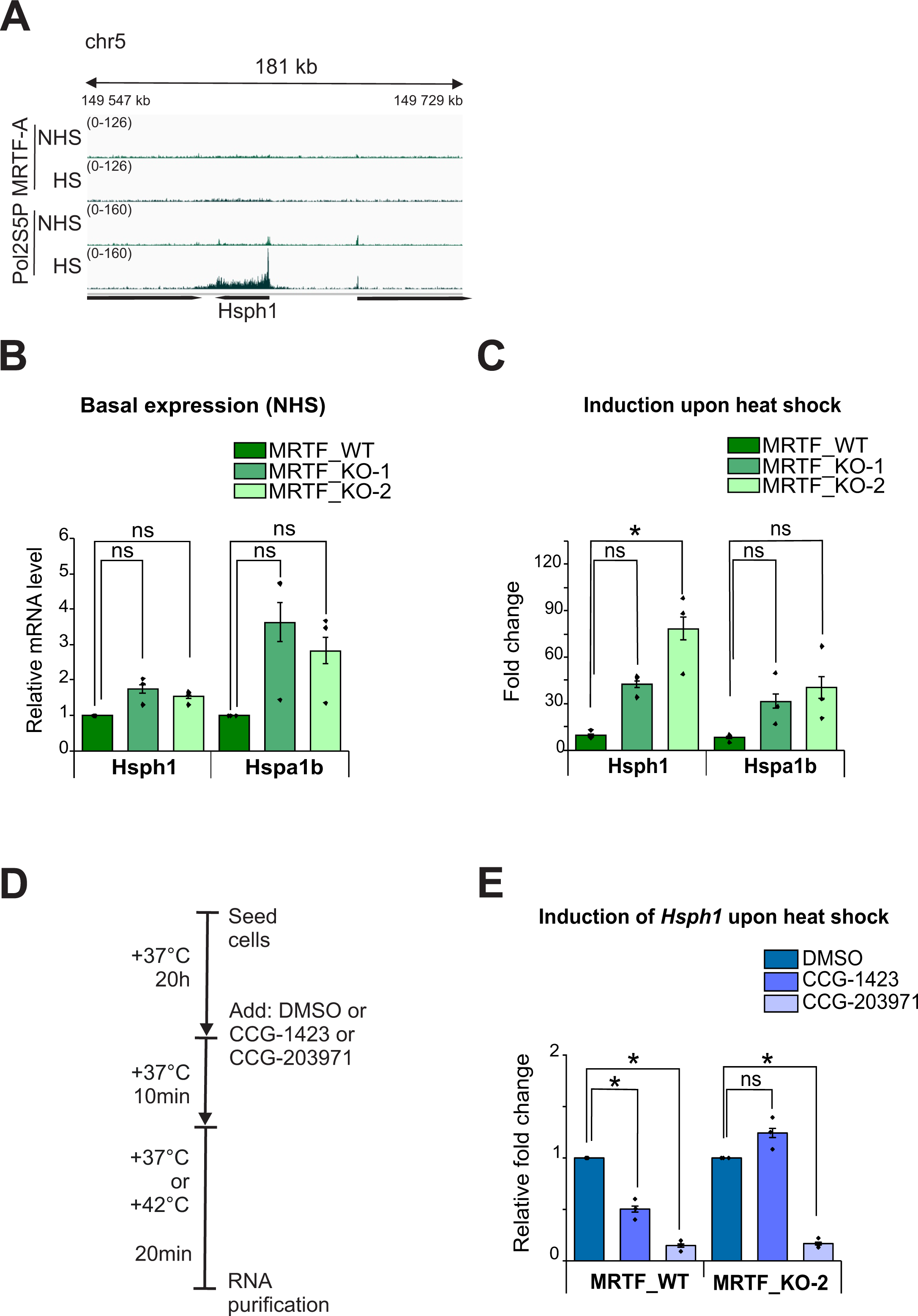
MRTFs are dispensable for *hsp* induction in mammalian cells, but CCG-derived compounds inhibit *hsp* induction also in the absence of MRTFs **A.** Normalized [Reads Per Kilobase per Million mapped reads (RPKM)] coverage of MRTF-A and RNA polymerase II phosphorylated on serine 5 (Pol2S5P) on heat shock responding gene *Hsph1* in mouse NIH 3T3 cells in non-heat shock (NHS) and heat shock (HS) conditions from ChIP-seq. **B.** Expression levels of *hsp* genes in the absence of MRTFs. Baseline expression of *Hsph1* and *Hspa1b* mRNAs in wild type (MRTF_WT) and two MRTF-KO (MRTF_KO-1 and MRTF_KO-2) mouse NIH 3T3 cell lines. Data is from three biological replicates, normalized to WT values, bars represent the mean value, individual data points are shown and error bars are standard errors of the mean (s.e.m). Statistical significances are measured with student’s one-sample t-test (*; P< 0.025 with Bonferroni correction). P-values: MRTF_WT vs MRTF_KO-1 for *Hsph1* is 0.08 MRTF_WT vs MRTF_KO-2 for *Hsph1* is 0.043. MRTF_WT vs MRTF_KO-1 for *Hspa1b* is 0.14, MRTF_WT vs MRTF_KO-2 for *Hspa1b* is 0.13. **C.** Induction of *hsp* gene transcription in the absence of MRTFs. Fold change in transcriptional induction of *hsp* genes upon 20 min heat shock. Data is from three individual experiments and shown as in B. Statistical significances are measured with one-way ANOVA followed by Tukey’s multiple comparison test (*; P< 0.05). P-values: MRTF_WT vs MRTF_KO-1 for *Hsph1* is 0.09, MRTF_WT vs MRTF_KO-2 for *Hsph1* is 0.004, MRTF_WT vs MRTF_KO-1 for *Hspa1b* is 0.27 and MRTF_WT vs MRTF_KO-2 for *Hspa1b* is 0.13. **D.** Schematic representation of the experimental design in mouse NIH 3T3 cells for data shown in E. **E.** Effect of CCG-derived inhibitors on *hsp* gene induction in the presence and absence of MRTFs. Data is the fold change in *Hsph1* mRNA induction upon 20 min of heat shock, from three individual experiments and shown as in B. Statistical significances are measured with student’s one-sample t-test (*; P< 0.025 with Bonferroni correction). P-values: CCG-1423 vs DMSO in MRTF_WT is 0.013, CCG-203971 vs DMSO in MRTF_WT is 0.0013, CCG-1423 vs DMSO in MRTF_KO-2 is 0.113 and CCG-203971 vs DMSO in MRTF_KO-2 is 0.0011. See also Supplementary Figure 2 for additional data.

To study if MRTFs are nevertheless required for *hsp* gene expression in mammalian cells, we used CRISPR/Cas9 to generate MRTF double knockout NIH3T3 (MRTF-KO) cells, which do not express either MRTF-A or MRTF-B. Clones were sequenced for the respective mutations and Western blotting was used to confirm the absence of protein products (Fig S2C). Moreover, these clones displayed significantly impaired induction of canonical MRTF/SRF target genes (Fig S2D). We then studied expression of *Hspa1b* and *Hsph1* genes in the presence and absence of HS (20 min, 42°C). Basal expression of *Hsph1* was not affected in either MRTF-KO clones examined compared to wild type (WT) control cells, while Hsph1 expression was modestly increased in one of the MRTF-KO clones (Fig 2B). Both *hsp* genes examined were efficiently activated in the MRTF-KO cells upon HS, *Hsph1* even more efficiently than in WT cells (Fig 2C). We then studied the effects of CCG-1423-derived inhibitors on *hsp* induction (Fig 2D). In WT cells, both CCG-1423 and CCG-203971 reduced HS-induced activation of *Hsph1* (Fig 2E). Intriguingly, CCG-203971 significantly inhibited *Hsph1* activation also in MRTF-KO cells (Fig 2E). This indicates that the effect of at least CCG-203971 on *Hsp* activation is not dependent on the presence of MRTFs. Combined, our results imply that MRTF-driven *hsp* induction is not conserved from fly to mammalian cells, but the effect of CCG-derived inhibitors is.

### CCG-derived inhibitors reduce transcription in fly and mammalian and cells

Since CCG-203971 inhibited *hsp* gene activation in the absence of MRTFs (Fig 2E), we wondered whether the CCG-derived compounds may influence gene transcription beyond the MRTF/SRF pathway. To test the specificity of the CCG-derived inhibitors we used SRF-reporter assays, based on firefly luciferase gene transcription regulated by serum response elements (SREs) (Geneste et al., 2002) together with HSV-thymidine kinase promoter controlling renilla luciferase expression as an internal normalisation control. When normalising the firefly luciferase to the respective renilla luciferase activity, both CCG-derived inhibitors seemed to reduce the SRF reporter activity in serum starved (Fig S3A) and serum stimulated conditions (Fig S3B), as reported before (Evelyn et al., 2007). However, when examining the raw luciferase activities, we noticed that CCG-derived inhibitors significantly reduced both the firefly and renilla luciferase activity under serum starved and stimulated conditions (Fig S3C-F). Since the CCG-derived inhibitors influence the transcription from also the HSV-thymidine kinase promoter-controlled vector, their effects seem not to be confined to SRF-mediated transcription.

To further explore how CCG-1423-derived inhibitors affect gene expression, we tested another inducible transcription system, namely ecdysone-induced transcription of *Eip74ef.* Treatment of the S2R+ fly cells with CCG-1423 did not influence basal expression of *Eip74ef* (Fig 3A), but significantly inhibited its transcriptional induction upon 1 h of 20H-ecd stimulation (Fig 3B). Actinomycin D (ActD), a well-established transcription inhibitor, also decreased hormone-induced expression of *Eip74ef* (Fig 3B). To measure global changes in RNA synthesis, we employed 5-ethynyl uridine (5-EU) incorporation into live fly S2R+ (Fig 3C) and mammalian NIH3T3 (Fig 3D) cells. Remarkably, both CCG-1423 and CCG-203971 significantly reduced 5-EU incorporation in fly (Fig 3C) and mammalian cells (Fig 3D). In S2R+ cells, the inhibition of transcription with CCG-1423-derived compounds was as strong as with ActD (Fig 3C). We conclude that CCG-1423-derived inhibitors globally inhibit RNA synthesis in fly and mammalian cells.

**Figure 3:**
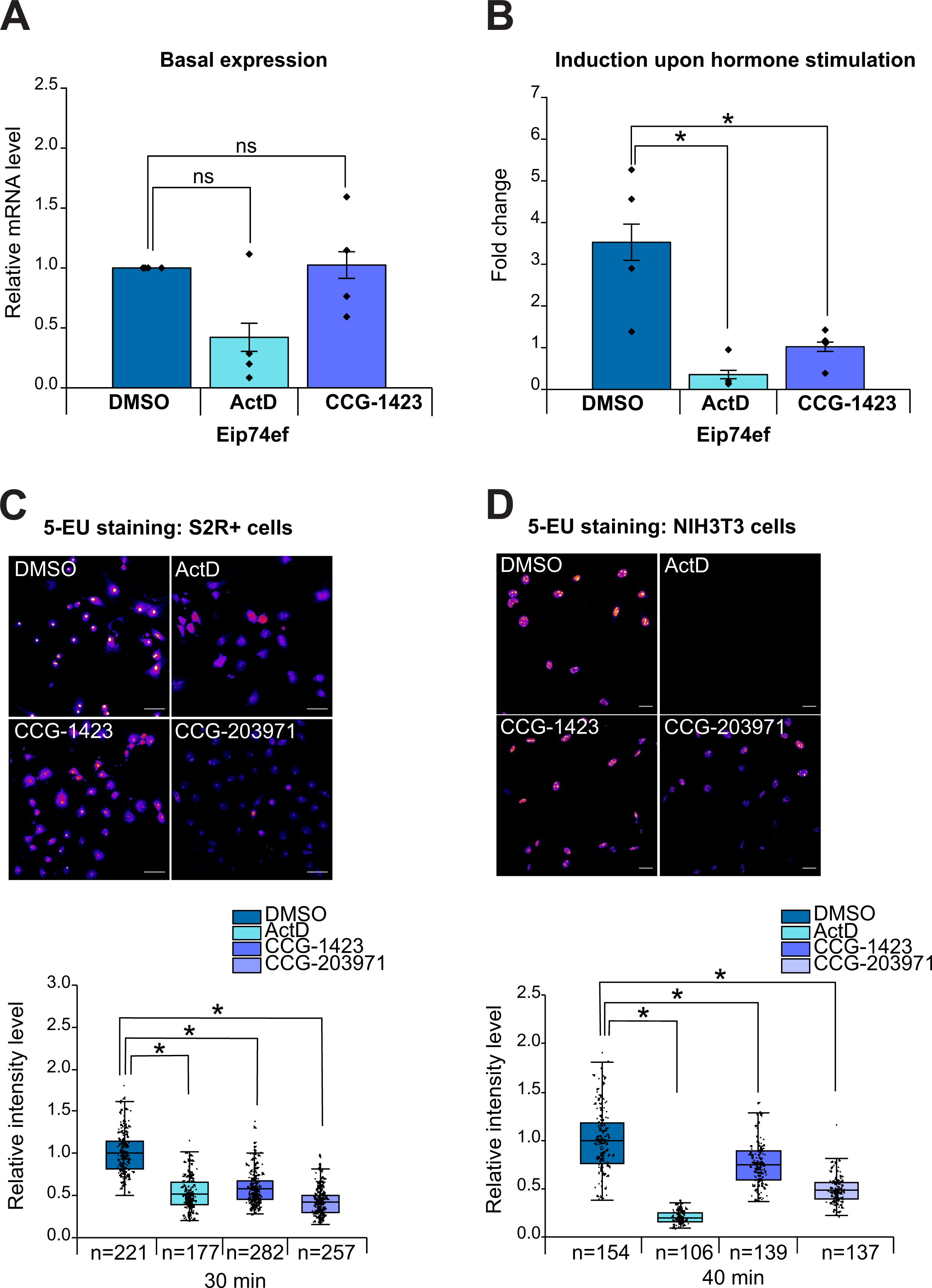
CCG-1423-derived inhibitors act as general transcription inhibitors **A.** Effect of CCG-1423 and Actinomycin D (ActD) on baseline expression of Ecdysone responsive gene *Eip74ef* in S2R+ fly cells. Data is normalized to DMSO treated sample, from four biological replicates, bars represent the mean value, individual data points are shown and error bars are standard errors of the mean (s.e.m). Statistical significances are measured with student’s one-sample t-test (*; P< 0.025 with Bonferroni correction). P-values: DMSO vs ActD is 0.09 and DMSO vs CCG-1423 is 0.92. **B.** Effect of CCG-1423 and Actinomycin D (ActD) on transcriptional induction of Eip74ef upon 1 h of 20H-ecdysone stimulation. Data is fold changes in *Eip74ef* mRNA levels, from four biological replicates and shown as in A. Statistical significances are measured with one-way ANOVA followed by Tukey’s multiple comparison test (*; P< 0.025 with Bonferroni correction). P-values: DMSO vs ActD is 0.0006 and DMSO vs CCG-1423 is 0.021. **C.** Effect of CCG-derived compounds and ActD on nascent RNA production as measured by 5 ethynyl uridine (5-EU) incorporation in fly S2R+ cells. Top: Fluorescence microscopy images of representative cells, scale bar 30 microns. Bottom: Quantification of mean intensities of 5-EU staining in the indicated conditions from three biological replicates is shown as a 25th and 75th percentiles box plot, where the midpoint indicates the median value of EU intensities, the box represents mean and whiskers represent data range within 1.5IQR. Measurements from single cells shown as data points, with the total number of quantified cells indicated below the graph. Statistical significance with Mann-Whitney U-test (*; P< 0.017 with Bonferroni correction). P-values: DMSO vs ActD is < 0.0001, DMSO vs CCG-1423 is <0.0001, DMSO vs CCG-203971 is <0.0001. **D.** Effect of CCG-derived compounds and ActD on nascent RNA production as measured by 5 ethynyl uridine (5-EU) incorporation in mouse NIH 3T3 cells. Data is shown as in C. Statistical significance with Mann-Whitney U-test (*; P< 0.017 with Bonferroni correction). P-values: DMSO vs ActD is <0.0001, DMSO vs CCG-1423 is <0.0001, DMSO vs CCG-203971 is <0.0001.

### CCG-derived inhibitors change the genome-wide distribution of RNA polymerase II

To understand how CCG-derived inhibitors influence transcription, we first used ChIP-seq to study the occupancy of RNA polymerase II in its serine-5 phosphorylated form (Pol II S5P) on chromatin in S2R+ cells with or without HS. Here we focused on CCG-1423, which had a more pronounced effect on *hsp* induction than CCG-203971 in these cells (Fig 1D). In non-HS conditions, CCG-1423 caused an accumulation of Pol II S5P on *hsp* gene bodies (defined as sequence +250bp from the transcription start site TSS to the cleavage and polyadenylation site CPS; see also materials and methods) (Fig 4A-B,D). Conversely, upon HS, CCG-1423-treated cells displayed substantially reduced Pol II S5P occupancy throughout the *hsp* genes as compared to DMSO treated cells (Fig 4A,C-D). The reduction in Pol II S5P occupancy in CCG-1423-treated cells was especially evident in the termination window, a region downstream of the CPS site (Fig 4A,C,E). On *hsp* genes, both Pol II S5P occupancy and transcriptional output measured as mRNA expression thus displayed similar responses to CCG-1423 treatment: increased Pol II density and mRNA levels in non-HS, but decreased inducibility upon HS. Beyond *hsp* genes, analysis of the 4981 expressed genes in S2R+ cells did not reveal significant, consistent differences between Pol II S5P levels on the gene body (Fig 5A, C) or the TSS (Fig 5A,B,D) in either non-HS or HS conditions. However, in non-HS conditions, CCG-1423 seemed to shift the pause-site of Pol II closer to the TSS (Fig 5B). In addition, the levels of Pol II S5P were significantly reduced after CPS in CCG-1423 treated cells in non-HS conditions (Fig 5A,E). These results indicate that in fly cells, CCG-1423 globally influences the genomic distribution of Pol II, and the effect is most pronounced at the end of the genes, suggesting changes in either Pol II processivity or transcript cleavage.

**Figure 4:**
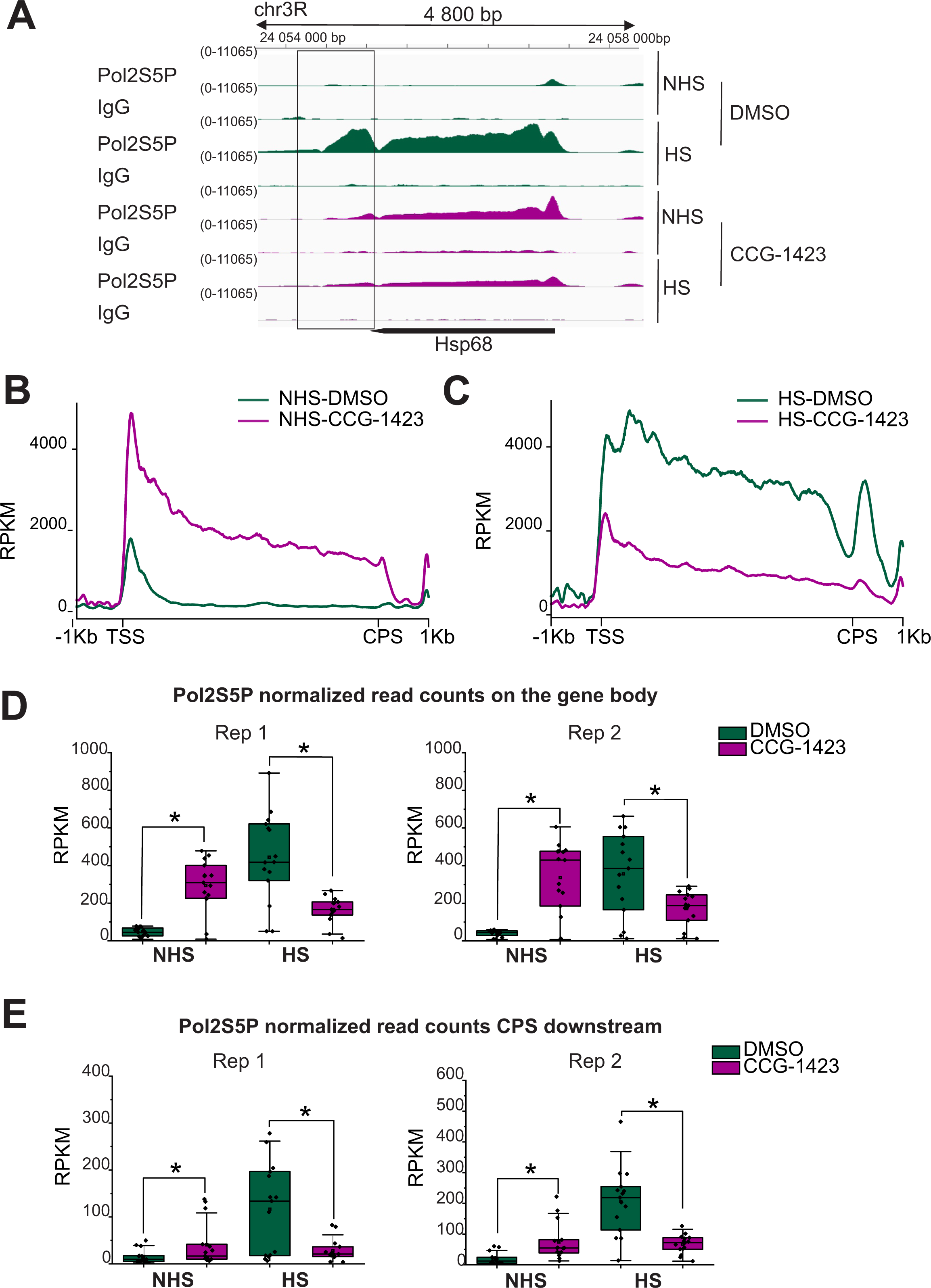
CCG-1423 influences Pol II binding to *hsp* genes both in non-heat shock conditions and upon heat shock **A.** Normalized (RPKM) coverage of RNA polymerase II phosphorylated on serine 5 (Pol2S5P) and IgG on heat shock responding gene *Hsp68* in S2R+ fly cells in DMSO and CCG-1423-treated cells in non heat shock (NHS) and heat shock (HS) conditions from ChIP-seq. Experimental set up as in Figure 1B. Box marks the 1Kb area downstream of the cleavage and polyadenylation site (CPS) used for quantification in E. **B.** Metaprofiles with average normalized fragment counts scaled to 5 kb gene size of Pol2S5P coverage on 15 heat shock responding genes in DMSO (green) and CCG-1423 (purple) treated cells in non-heat shock conditions. **C.** Metaprofiles of Pol2S5P coverage on the 15 heat shock responding genes in DMSO and CCG-1423 treated cells upon heat shock. **D.** Quantification of Pol2S5P on the gene body of 15 heat shock responding genes as normalized read counts (RPKM) in DMSO (dark green) and CCG-1423 treated (magenta) cells in non-heat shock (NHS) and heat shocked (HS) conditions in two replicate experiments. Data is shown as a 25th and 75th percentiles box plot, where the horizontal line represents the median, small box represents mean and whiskers represent data range within 1.5IQR. Statistical significance with Mann-Whitney U test (*; P< 0.05). P-values: replicate 1 NHS is 0.0002 and HS is 0.019; replicate 2 NHS is 0.0001 and HS is 0.0005. **E.** Quantification of Pol2S5P 1 kb after CPS of 15 heat shock responding genes in DMSO and CCG-1423 treated cells in non-heat shock (NHS) and heat shocked (HS) conditions in two replicate experiments. Data is shown as in D. Statistical significance with Mann-Whitney U test (*; P< 0.05). P-values: replicate 1 NHS is 0.04 and HS is 0.03, replicate 2 NHS is 0.001 and HS is 0.0002. See supplementary figure 4 for the respective data from mouse cells.

**Figure 5:**
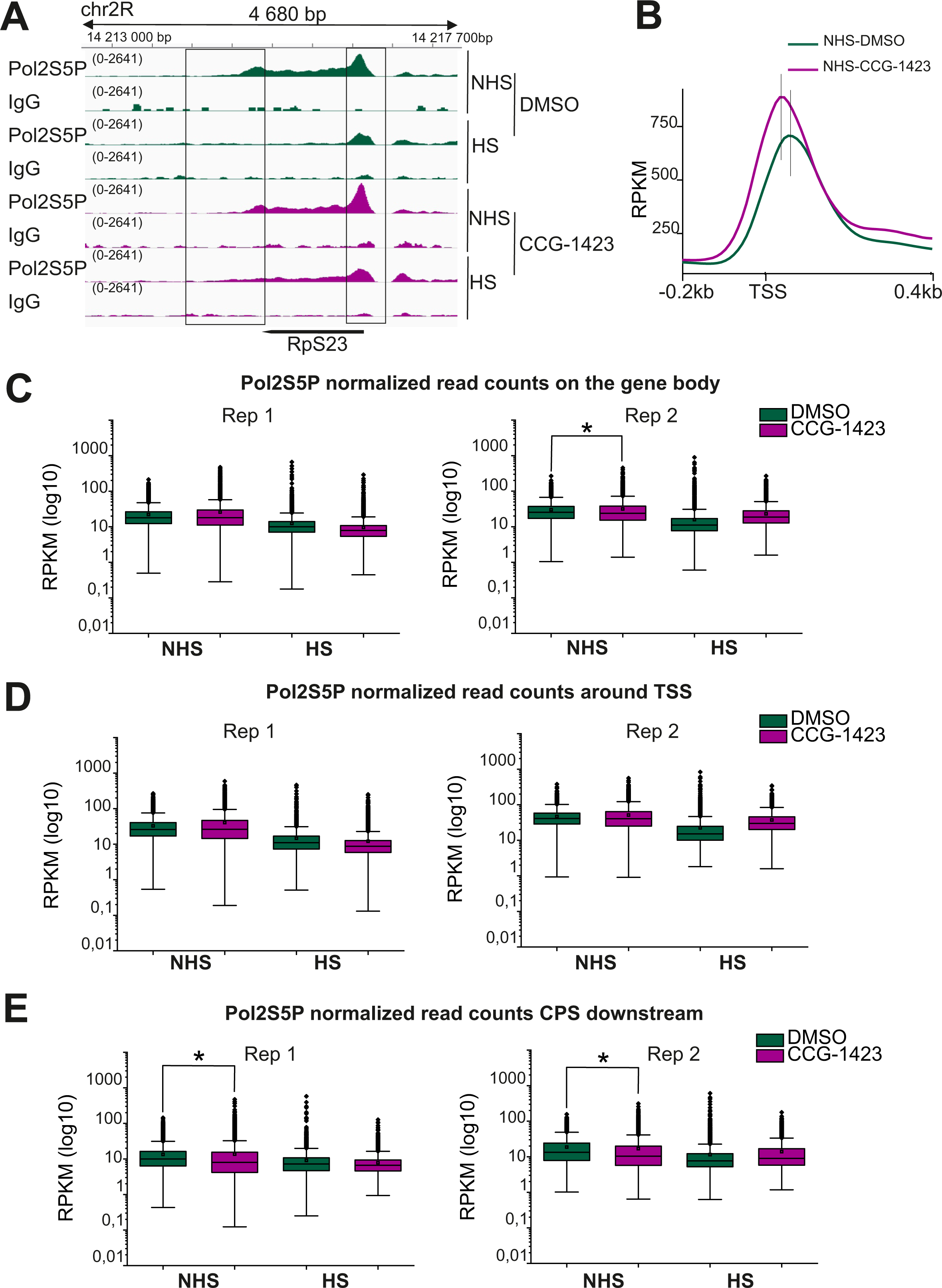
CCG-1423 influences genome-wide occupancy of RNA polymerase II at the end of genes **A.** Normalized (RPKM) coverage of RNA polymerase II phosphorylated on serine 5 (Pol2S5P) and IgG on *Rps23* gene in S2R+ fly cells in DMSO and CCG-1423-treated cells in non-heat shock (NHS) and heat shock (HS) conditions from ChIP-seq. Experimental set up as in Figure 1B. Regions of transcription start site (TSS) (from −250bp to +250bp) and 1Kb area downstream of the cps cleavage and polyadenylation site (CPS) used for quantification in D and E are indicated with a box. **B.** Coverage of Pol2S5P around the TSS region of 4981 expressed genes in DMSO and CCG-1423 treated cells in non-heat shock conditions shown as metaprofiles with average normalized fragment counts. CCG-1423 shifts the pause site (indicated with a vertical line) closer to TSS. **C.** Quantification of Pol2S5P on the gene body of the 4981 expressed genes as normalized read counts (RPKM) in DMSO (dark green) and CCG-1423 treated (magenta) cells in non-heat shock (NHS) and heat shocked (HS) conditions in two replicate experiments. Data is shown as a 25th and 75th percentiles box plot, where the horizontal line represents the median, small box represents mean and whiskers represent data range within 1.5IQR. Data is shown as a 25th and 75th percentiles box plot, where the horizontal line represents the median, small box represents mean and whiskers represent data range within 1.5IQR. Statistical significance with Mann-Whitney U test (*; P< 0.05). P-value for NH DMSO vs CCG-1423: replicate 2 is 0.0003. **D.** Quantification of Pol2S5P around TSS of the 4981 expressed genes shown as in C. **E.** Quantification of Pol2S5P downstream of CPS of the 4981 expressed genes shown as in C. Statistical significance with Mann-Whitney U test (*; P< 0.05). P-values for NH DMSO vs CCG-1423: replicate 1 is <1E-100 and replicate 2 is <1E-100 See supplementary figure 5 for the respective data from mouse cells.

To study, if the effects of CCG-derived inhibitors on Pol II are conserved in mammalian cells, we performed ChIP-seq analysis of Pol II S5P in NIH 3T3 cells with CCG-203971 (Figure S4). Analysis of 15 *hsp* genes did not reveal any statistically significant effects of CCG-203971 in either non-HS or HS condition, likely due to the low number of *hsp* genes analysed. However, the trend especially regarding the decreased Pol II S5P signal after the CPS upon CCG-203971 treatment and HS (Fig S4D) was similar as observed for fly cells (Fig 4F). Thus, in line with qPCR analysis of *hsp* transcript levels (Fig 2E), CCG-derived compounds had a smaller effect on Pol II in mammalian cells than in fly cells. Global analysis of the 11468 expressed genes in NIH 3T3 cells revealed significant increase in Pol II S5P signal near the TSSs in CCG-203971 treated cells in non-HS conditions (Fig S5A,C), although the signal on the gene bodies (Fig S5B) or after CPS (Fig S5D) were not significantly and consistently affected. Upon HS, no significant differences were detected (Fig S5A-D). We conclude that also in mammalian cells, CCG-derived inhibitors globally influence Pol II function. However, the effects of CCG-derived inhibitors on RNA synthesis manifest differently in fly and mouse cells, likely due to subtle differences in overall transcriptional regulation between different organisms.

### CCG-1423 prevents transcriptional reprogramming upon HS

To gain a mechanistic understanding on how CCG-1423-derived inhibitors influence transcription, we performed precision run-on sequencing (PRO-seq) (Kwak et al., 2013; Mahat et al., 2016a) in fly S2R+ cells. We used whole-genome spike-in normalisation to enable comparison of transcription activity between conditions and collected each sample in two biological replicates that showed high correlation (Rho > 0.93; Fig S6A). Taking advantage of the strand specificity and nucleotide-resolution mapping of engaged Pol II, we quantified Pol II density on the coding strand of each gene to reveal differentially expressed genes (p-value<0.05) upon HS and CCG-1423 treatments (Figure 6A, Supplementary Table S1). In DMSO treated control cells, HS triggered an induction of 569 and repression of 5520 genes (Figure 6A), which agrees well with previous studies on HS-induced transcriptional responses in fly S2 cells (Duarte et al., 2016). As expected, gene ontology analysis revealed that the heat-induced genes were enriched with functions related to stress responses and protein refolding (Fig S6B, Supplementary Table S2), whereas the functions enriched among the downregulated genes were comprised of cell division, translation, mRNA processing and protein transport (Fig S6C, Supplementary Table S2). In basal, non-HS conditions, CCG-1423 treatment resulted in upregulation of 16 genes (Fig 6A), 13 of which (Fig S6B) belong to the classical heat-inducible genes (reviewed by Vihervaara et al. 2018). Intriguingly, only this same subset, together with 5 additional genes, were upregulated in the CCG-1423-treated HS sample, when compared to DMSO treated non-HS sample (Fig 6A,B, Fig S6B). This indicates that CCG-1423 prevents the induction of the vast majority of HS-inducible genes. Analysis of the engaged Pol II on *hsp* genes revealed a similar trend as observed by ChIP-seq (Fig 4): CCG-1423 treatment resulted in accumulation of transcribing polymerase across *hsp* genes in non-HS conditions, but it prevented further increase of transcribing polymerase on these genes upon HS (Fig 6C-D), including the termination windows after CPS (Figure 6E). Indeed, extending the analysis to all heat-induced genes (n=212; >2500 nt in length) revealed the genome-wide inability to activate transcription upon HS in the presence of CCG-1423 (Fig 6F).

**Figure 6.**
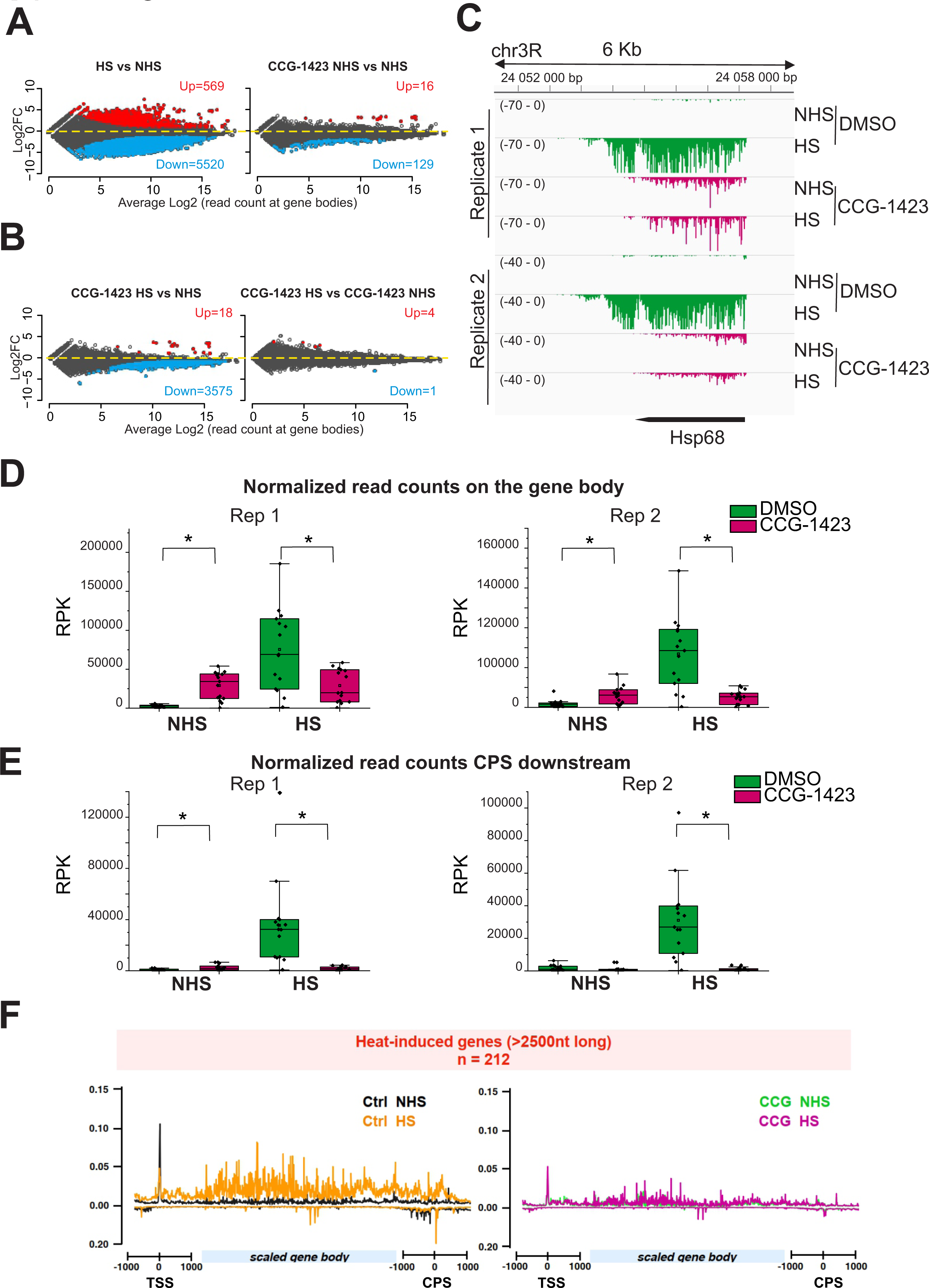
CCG-1423 severely dampens heat-induced transcriptional reprogramming **A.** Differential gene transcription upon heat shock (HS) (left) and CCG-1423 treatment (right) in fly S2R+ cells, as compared to non-heat shocked (NHS) DMSO-treated control cells. Up (red) and down (blue) denote statistically significant (p<0,05) increase or decrease in RNA polymerase II density at gene bodies from PRO-seq. **B.** Differential gene transcription upon heat shock (HS) in the presence of CCG-1423 in S2R+ fly cells, as compared to non-heat shocked (NHS) DMSO-treated control cells (left), or to non-heat shocked (NHS) but CCG-1423-treated cells (right). Up (red) and down (blue) denote statistically significant (p<0,05) increase or decrease in RNA polymerase II density at gene bodies from PRO-seq. **B.** Venn diagram showing the overlap of genes activated upon HS in DMSO and CCG-1423 treated cells. Gene names indicated for the genes activated by HS in both conditions and enriched gene ontologies for genes activated by HS exclusively in DMSO treated cells. **C.** Density of engaged Pol II (PRO-seq) at heat-induced *Hsp68* gene in S2R+ fly cells in non-heat shock (NHS) and heat shock (HS) conditions in DMSO or CCG-1423 treated cells. **D.** Quantification of spike-in normalized PRO-seq read counts (RPK) on gene bodies of the 15 heat shock responding genes in DMSO and CCG-1423 treated cells in non-heat shock (NHS) and heat shocked (HS) conditions in two replicate experiments. Data is shown as a 25th and 75th percentiles box plot, where the horizontal line represents the median, small box represents mean and whiskers represent data range within 1.5IQR. Each data point is indicated as a dot and both replicates are shown. Statistical significance with a Mann-Whitney U test (*; P< 0.05). P-values: replicate 1 NHS is 0.00005 and HS is 0.011, replicate 2 NHS is 0.009 and HS is 0.0004. **E.** Quantification of spike-in normalized PRO-seq read counts (RPK) downstream of transcription end site (CPS) at the 15 heat shock responding genes as shown in B. Statistical significance with a Mann-Whitney U test (*; P< 0.05). P-values: replicate 1 NHS is 0.004 and HS is 0.00003, replicate 2 NHS is NS and HS is 0.00003. **F.** Average density of engaged Pol II across heat-induced genes (>2500nt long, n=212) in DMSO and CCG-1423 treated cells in non-heat shock (NHS) and heat shocked (HS) conditions.

Upon HS, fewer genes were down-regulated in the CCG-1423 treated cells (n=3575) compared to DMSO treated cells (n=5520), and the overall level of transcriptional repression was blunted (Fig 6A,B). The HS-repressed genes in DMSO and CCG-1423 treated cells displayed 81% overlap (n=2889), suggesting a genome-wide change in the ability to repress transcription (Fig S6C). Interestingly, the genes that were downregulated specifically by HS, and not in the CCG-1423 treated HS sample, were enriched in processes related to cytoplasmic translation (Fig S6C). However, genes specifically repressed in the CCG-1423 treated, but not in DMSO-treated HS samples, were more enriched in processes related to nuclear transcription (Fig S6C). As an example of a gene that is downregulated by HS, we analysed the transcribing polymerase on the *Rps23* gene. As expected, in DMSO treated cells HS induced a marked reduction in the transcribing polymerase along the whole gene (Fig 7A, FigS7A). However, in CCG-1423 treated cells, this reduction was clearly less pronounced (Fig 7A, FigS7A), indicating that CCG-1423 dampens HS-induced transcriptional repression. In support of this, analysis of all expressed genes (n=11468) upon HS revealed a significant increase in the engaged polymerase in CCG-1423-treated cells compared to DMSO treated cells, both at the gene body (Fig 7B), and termination window (Fig 7C). Notably, ChIP-seq failed to detect these effects (Fig 5), which could indicate that the ability of Pol II to catalyse the addition of ribonucleotides to the nascent RNA chain, required for PRO-seq signal, is reduced in the presence of CCG-1423. Finally, metaprofiles of the transcribing polymerase specifically on heat-repressed genes (n=2039; >2500 nt in length) were essentially identical in CCG-1423 treated samples with and without HS (Fig 7D), indicating the drastically reduced ability to respond to HS. Combined, these results reveal that CCG-1423 severely dampens the overall transcriptional reprogramming upon HS by inhibiting both the transcriptional activation and genome-wide repression. Taken together, these results uncover that CCG-derived compounds manifest a genome-wide effect on Pol II and transcriptional responsiveness and suggest inhibition of Pol II processivity (Fig 7E).

**Figure 7.**
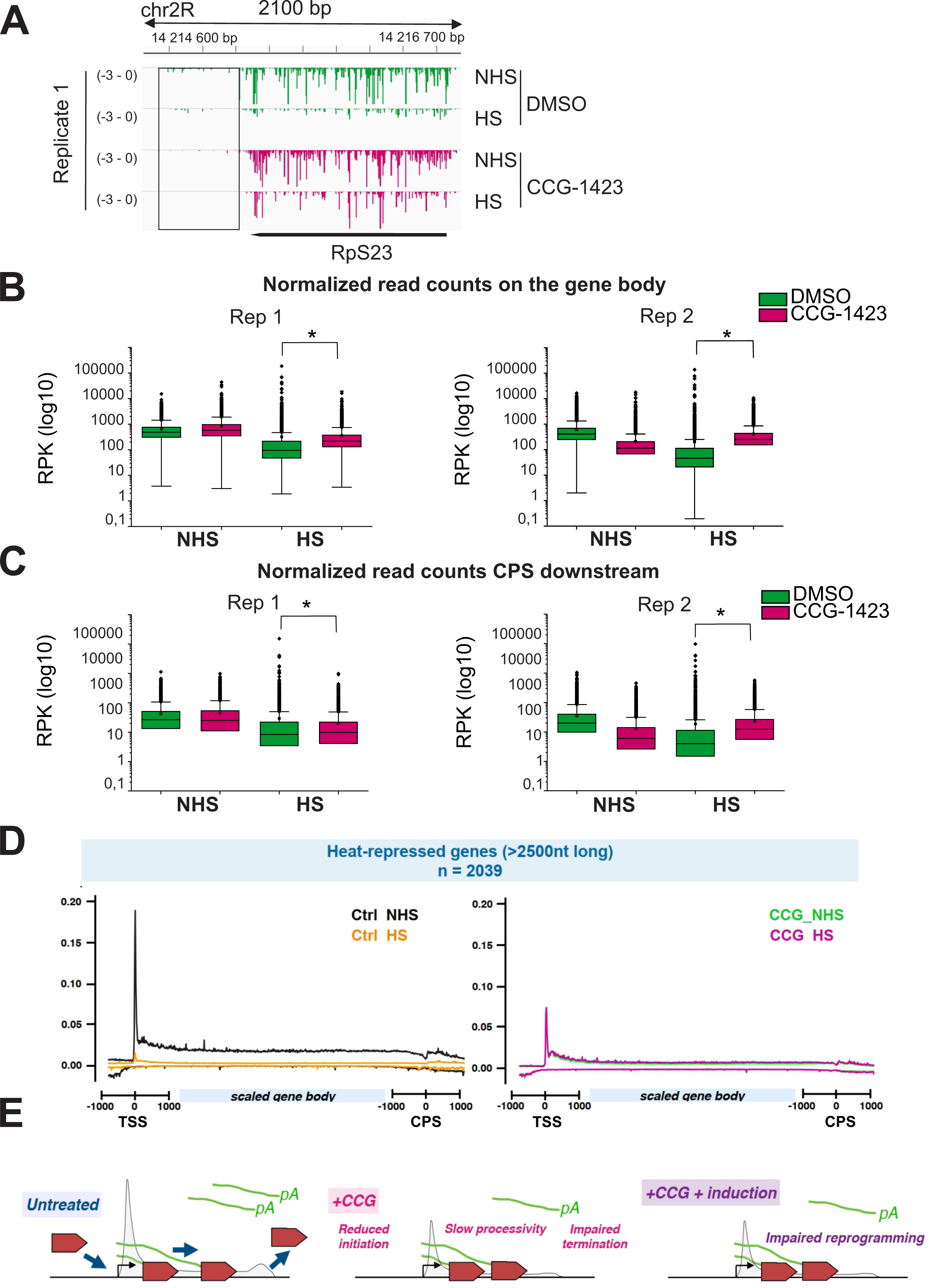
CCG-1423 reduces transcriptional repression upon heat shock **A.** Density of engaged Pol II (PRO-seq) at *Rps23* gene in S2R+ fly cells in non-heat shock (NHS) and heat shock (HS) conditions in DMSO or CCG-1423 treated cells. **B.** Quantification of spike-in normalized read counts (RPK) on gene bodies of the 4981 expressed genes in DMSO and CCG-1423 treated cells in non-heat shock (NHS) and heat shocked (HS) conditions in two replicate experiments. Data is shown as a 25th and 75th percentiles box plot, where the horizontal line represents the median, small box represents mean and whiskers represent data range within 1.5IQR. Statistical significance with Mann-Whitney U test. (*; P< 0.05). P-values for NH DMSO vs CCG-1423: replicate 1 is 2E-63 and replicate 2 is 1E-00. **C.** Quantification of spike-in normalized read counts (RPK) downstream of the transcription end site (CPS) of the 4981 expressed genes in DMSO and CCG-1423 treated cells in non-heat shock (NHS) and heat shocked (HS) conditions shown as in B. Statistical significance with Mann-Whitney U test (*; P< 0.05). P-values for NH DMSO vs CCG-1423: replicate 1 is <0.0003 and replicate 2 is <2E-285. **D.** Average density of engaged Pol II across heat-repressed genes (>2500nt long, n=2039) in DMSO and CCG-1423 treated cells in non-heat shock (NHS) and heat shocked (HS) conditions. **E.** Model on how CCG-1423-derived compounds impair transcriptional responses.

## Discussion

Small molecule inhibitors are attractive research tools and even therapeutic reagents due to their ease of use, since they often do not require elaborate protocols for their delivery into cells or tissues. Nevertheless, understanding the specificity of the inhibitor, and mechanism of action, are keys for its successful and predictable use in any application. Here we report that CCG-1423-derived inhibitors, which have been extensively used as specific inhibitors of the MRTF/SRF transcription pathway, have a global effect on transcription. These results necessitate re-evaluation of the results obtained with these compounds, but also open new possibilities for their application in transcription research.

Our initial aim for utilising CCG-derived inhibitors was to study the effects of MRTFs on *hsp* gene transcription motivated by earlier studies from *Drosophila* ovaries (Salvany et al., 2014; Somogyi and Rorth, 2004). However, we find that in mammalian cells, *hsp* genes were efficiently activated in fibroblasts lacking MRTF-A and MRTF-B (Fig 2C), and we also failed to detect MRTF binding to *hsp* gene promoters in these cells (Fig 2A). It is therefore possible that the role of MRTF in *hsp* expression is fly-specific and not conserved in mammalian cells. It might even be that the role is restricted to fly ovaries, since at least in some *hsp* genes, separate regulatory elements seem to be required for heat-inducible and ovarian-specific expression (Cohen and Meselson, 1985). Development of new reagents, for example an antibody recognizing MAL-D, would be needed to explore this further.

CCG-1423-derived inhibitors have been widely used to inhibit transcriptional responses downstream of Rho-signalling via MRTF/SRF pathway, and in this capacity, to inhibit several cellular responses that depend on this pathway, including cancer cell migration and fibrosis (Haak et al., 2017; Lundquist et al., 2014; Minami et al., 2012; Sisson et al., 2015). Our results, obtained with several different methods, from luciferase-based reporter assays (Fig S3) to endogenous target gene expression (Fig 1,2) and 5-EU incorporation (Fig 3C-D), as well as ChIP-seq (Figs 4-5) and PRO-seq (Figs 6-7) demonstrate that the effects of CCG-derived inhibitors extend well beyond MRTF/SRF transcription pathway in both fly and mammalian cells. Interestingly, a microarray-based gene expression analysis of human PC-3 prostate cancer cell line had earlier revealed a relatively low overlap between MRTF-dependent and CCG-1423-regulated genes, leading to speculation that CCG-1423 could have a broader influence on transcription (Evelyn et al., 2016). Most published studies utilising CCG-1423-derived inhibitors use overnight incubation with the compounds, and therefore some of the observed effects are likely to be indirect. Nevertheless, we observed very robust inhibition of *hsp* gene transcription (Fig 1D) and 5-EU incorporation (Fig 3C-D) already upon 30 min treatment with CCG-1423-derived inhibitors. It can therefore be speculated that the effects reported in this study reflect more direct consequences of CCG-1423-mediated inhibition, perhaps via its primary target(s).

Several targets have been proposed for CCG-1423-derived inhibitors, ranging from MRTF itself (Hayashi et al., 2014) to actin regulators (Lundquist et al., 2014) and other transcription cofactors (Lisabeth et al., 2019). We find that CCG-203971 inhibits *hsp* gene transcription in the absence of MRTFs (Fig 2E), strongly arguing against the target being either MRTF or an actin regulator. Recently, pirin, an iron-binding transcription factor implicated in NF-Kb signaling, was identified as the molecular target of CCG-derived compounds with an affinity isolation-based target identification method using the CCG-222740 compound (Lisabeth et al., 2019). Nevertheless, we did not find any evidence that bisamide (CCT251236), which binds pirin with high affinity (Cheeseman et al., 2017) would inhibit either basal or induced expression of HS and serum-inducible target genes (Fig S8). Moreover, we have failed to identify an obvious pirin ortholog in *Drosophila melanogaster,* although CCG-1423 had a robust transcriptional effect in the fly S2R+ cells (Fig 1D, 3C). Further experiments are required to clarify the contribution of pirin in cellular responses to CCG-1423-derived compounds. Here, we focused our analysis on nascent transcription and Pol II progression, due to our initial interest in *hsp* gene expression. Of note, 5-EU (Fig 3C,D) is incorporated into all synthesized RNA, including ribosomal RNA, and qualitatively, the 5-EU signal emanating from nucleoli was also reduced in both fly and mammalian cells treated with CCG-derived compounds (Fig 3C,D). Moreover, PRO-seq revealed decreased transcribing Pol III on *tRNA* genes in CCG-1423-treated cells, as compared to DMSO-treatment (Fig S9). It is therefore possible that CCG-1423-derived compounds generally influence all RNA polymerases, necessitating further studies on the specificity of CCG-derived compounds on transcriptional regulation and cellular homeostasis. In this regard, it is also interesting to note that while CCG-1423 and CCG-203971 had differential effects on *hsp* transcription in fly and mammalian cells (Figs 1C-D, 2E), they both significantly reduced 5-EU incorporation in the cells tested here.

Regardless of the primary target, our ChIP- and PRO-seq experiments suggest that CCG-1423-derived compounds profoundly influence Pol II function. This manifests as a dampened transcriptional response to HS, reducing not only *hsp* gene activation, but also genome-wide transcription repression. The latter phenomenon was detectable only by PRO-seq (Fig 7), but not by ChIP-seq (Fig 5), suggesting defects in Pol II processivity, and attesting to the power of using these types of methods in combination to fully understand the regulation of transcription cycle (Wissink et al., 2019). Controlling the release of promoter-proximally paused Pol II is critical for transcriptional responses upon HS (Vihervaara et al., 2018). Indeed, in mammals, promoter-proximal Pol II pausing coordinates both heat-induced activation and repression, while in fly cells repression is primarily caused by reduced initiation. The promoter-proximal pause-release depends on positive transcription elongation factor b (P-TEFb), which could therefore be a plausible target for CCG-1423-derived inhibitors. However, inhibiting P-TEFb with flavopiridol, or more specifically by utilising analog-sensitive mutants of the kinase subunit of P-TEFb, Cdk9, leads to accumulation of Pol II to the promoter proximal pause site (Henriques et al., 2013; Laitem et al., 2015; Tellier et al., 2022), which we observe only in mammalian cells (Fig S5C), but not in fly cells (Fig 5D) upon CCG-treatment. Moreover, microarray analysis did not reveal significant overlap between CCG-1423 and Cdk9 inhibitor 5,6-dichloro-1-beta-D-ribofuranosylbenzimidazole (DRB)-regulated genes (Evelyn et al., 2016), indicating that these two compounds do not affect transcription similarly.

Intriguingly, in mammalian cells, both Cdk9 inhibition and HS have been shown to induce premature termination of transcription (Cugusi et al., 2022; Laitem et al., 2015; Tellier et al., 2022). Many factors involved with transition from transcription elongation to transcription termination are Cdk9 targets, and therefore loss of Cdk9 activity leads to premature termination of transcription and failure to polyadenylate the mRNA (Tellier et al., 2022). Upon HS, it has been postulated that inhibition of U1 telescripting, where U1 snRNP suppresses premature termination from intronic cryptic polyadenylation sites to ensure production of full-length transcripts, underlies premature termination and thereby transcription repression (Cugusi et al., 2022). Although U1 telescripting is conserved also in flies (Berg et al., 2012), its relevance to HS-induced transcriptional changes in this organism has not been documented to our knowledge. Our working hypothesis is that also CCG-1423 treatment causes premature termination of transcription by reducing Pol II processivity (Fig 7E). This could take place by affecting Pol II elongation rate and/or its coupling to co-transcriptional processes. Premature termination upon CCG-1423 treatment could explain why these compounds reduce Pol II occupancy especially at the end of the genes (Fig 4E, 5E, 6E, 7C). Elongation rate of Pol II has a major impact on co-transcriptional processes, such as splicing and transcription termination (reviewed in (Muniz et al., 2021)). We find that CCG-1423 treatment causes inefficient clearing of Pol II during HS-induced transcriptional repression (Fig 7), suggesting that CCG-1423 might slow down Pol II elongation. Indeed, we observe shifting of the pause site closer to TSS in CCG-1423 treated cells (Fig 5B), similarly as reported before for a slow Pol II mutant in *Drosophila* (Li et al., 2013).

The many phenotypic effects of CCG-derived compounds reported in the literature could result from the overall dampening of transcriptional responses to different signalling pathways. This includes the MRTF/SRF transcription pathway, which can be strongly activated by various signals, and therefore likely very susceptible to these inhibitors. In the future, it will be important to develop novel approaches to elucidate the direct target(s) of CCG-1423-compounds. This would facilitate their use in transcription research, since they seem to influence Pol II function by novel mechanism that is distinct from the most commonly used transcription inhibitors.

## Acknowledgements

We acknowledge Paula Maanselkä for excellent technical assistance throughout this project and Marko Crivaro for the help with statistics. Imaging was performed at the Light Microscopy Unit, sequencing at Biomedicum Functional Genomics Unit (FuGU), and fly work was supported by HiFLY, all supported by HiLIFE and Biocenter Finland. Flow cytometry was performed at Viikki Flow Cytometry, supported by HiLIFE. Part of sequencing was performed at Novogene. We thank Guido Posern (University of Halle) for providing the MRTF antibody. This work was supported by Sigrid Juselius, Cancer and Jane and Aatos Erkko foundations as well as Academy of Finland grants 338281 and 330254 to MKV, as well as Academy of Finland grant 324952, KTH and SciLifeLab (Fellowship) and Veteskapsrådet start grant 2021-02668 to AV.

## Materials and Methods

### Chemicals

Actinomycin D (A9415; Sigma Aldrich) was dissolved to an initial concentration of 5 mM in DMSO. CCG-1423 (sc-205241A; Santa Cruz Biotechnology) and CCG-203971 (SML1422; Sigma Aldrich) were dissolved to an initial concentration of 10 mM in DMSO. 20-hydroxyecdysone (20-E) (H5142; Sigma Aldrich) was resuspended to an initial stock concentration of 10 mM in ethanol.

### Cell lines and flies

*Drosophila* flies were maintained at +25°C. Fly strain mal-dΔ7 (#58418) were ordered from Bloomington Drosophila Stock Center.

Drosophila Schneider cell line 2 (S2R+) was cultured in Schneider’s Drosophila media (21720024, Gibco) supplemented with 10 % heat inactivated Fetal Bovine Serum (HI FBS) (10500064; Gibco) and 1x antibiotic-antimycotic drug (100 units/mL of penicillin, 100 µg/mL of streptomycin, and 0.25 µg/mL of Amphotericin B) (15240062; Gibco). Cells were cultured at a 25°C incubator in T75 flasks and passaged every 3 to 4 days.

Mammalian cell line NIH 3T3, MRTF-WT and MRTF knockout in NIH 3T3 (MRTF-KO-1 and MRTF-KO-2) cells were cultured in DMEM (BE12-614Q, Lonza) supplemented with 10 % Fetal Bovine Serum (FBS) (10270-106; ThermoFisher Scientific), 100 units/mL of Penicillin and 100 µg/mL Streptomycin (15140-122; ThermoFisher Scientific) and 1x of L-glutamine (35050061; ThermoFisher Scientific) and maintained in humidified 95% air by 5% CO_2_ incubator at + 37°C.

### Genome editing in mouse NIH 3T3

Generating MRTF knockout cell lines (MRTF-KO-1 and MRTF-KO-2) using CRISPR (clustered regularly interspaced short palindromic repeats) in complex with Cas9 (CRISPR associated protein 9) protein technology was performed according to previously published protocol (Mali et al., 2013).

The following 23 bp guide RNAs (gRNAs) in the coding region of MRTFA (exon5) and MRTFB (exon7) were designed using CHOPCHOP (Labun et al., 2019), CRISPOR (Haeussler et al., 2016), and GPP sgRNA Designer.

MRTF_A+B: TTCTTCTCCACCAGCTCCATGGG

Following oligoes were annealed and were incorporated into the gRNA cloning vector pSpCas9(BB)-2A-GFP, also known as PX458.

MRTF_A+B_Fwd: CACCGTTCTTCTCCACCAGCTCCAT

MRTF_A+B_Rev: AAACATGGAGCTGGTGGAGAAGAAC

After sequence verification, empty vector and vector containing insertions with MRTF_A+B gDNAs were transfected into the NIH 3T3 cells using JetPRIME transfection reagent according to the manufacturer’s protocol. The transfected NIH 3T3 cells were sorted using fluorescence-activated cell sorting (FACSAria II), which was based on the GFP signal of the cells. Cells were plated in a low density to grow single colonies. After several single-cell colonies had reached confluence they were split and levels of MRTF-A and MRTF-B were verified with Western Blotting using antibodies (Table 1) as described above (Fig S2C). In several clones MRTF-A and MRTF-B proteins were not detected and presence of mutation were verified with NGS of genomic DNA using MiSeq Nano (Biomedicum Functional Genomics Unit (FuGU)). Clone numbers 7-1 and 7-6 were chosen for experiments (MRTF_KO-1 and MRTF_KO-2 respectively). Clone number 1 from the empty vector transfection was chosen for experiments as control (MRTF_WT).

**Table 1.** List of antibodies used in the study

| Name | Catalogue; Manufacturer | Application |
| --- | --- | --- |
| GAPDH | GA1R; ThermoFisher Scientific | Western blotting |
| Histone 4 (H4) | SAB4500311; Sigma Aldrich | Immunofluorescent staining |
| MRTF-A | Home-made rabbit antiserum provided by G.Posern (Martin Luther University Halle-Wittenberg) | ChIP-seq |
| MRTF-A | #14760; Cell Signaling Technology | Western blotting |
| MRTF-B | #14613; Cell Signaling Technology | Western blotting |
| RNA polymerase II CTD repeat YSPTSPS (phospho S5) | ab5408; Abcam | ChIP-seq |
| Goat anti-Mouse IgG (H+L) Cross-Adsorbed Secondary Antibody, HRP | G-21040; Invitrogen/ThermoFisher Scientific | Western blotting |
| Goat anti-Rabbit IgG (H+L) Secondary Antibody, HRP | G-21234; Invitrogen/ThermoFisher Scientific | Western blotting |
| Goat anti-Rabbit IgG Highly Cross-Adsorbed Secondary Antibody, Alexa Fluor™ Plus 647 | A32733; Invitrogen | Immunofluorescent staining |
| Normal mouse IgG | Sc-2025; SantaCruz Biotechnology | ChIP-seq |

### dsRNA synthesis and RNA interference (RNAi)

Double stranded RNA (dsRNA) for target gene silencing was designed with SnapDragon design dsRNAs for fly cell RNAi. dsRNA was synthesised using MEGAscript® T7 Kit using manufacturer’s protocol (AMB13345; Invitrogen/ThermoFisher Scientific) and purified with NucleoSpin RNA kit (740955; Macherey-Nagel).

For RNAi, S2R+ cells were seeded at a density of 1 million cells/mL in serum free Schneider’s media in a six wells plate (1 mL per well). 4 ug dsRNA was added slowly to the media and allowed to incubate at 25°C for 1 hour. After the incubation period, 1 ml of Schneider’s media supplemented with 10 % HI FBS was added to each well. Cells were allowed to grow for 4 days at 25°C, then subjected to either heat shock at 37°C waterbath for 20 mins or non heat shock at 25°C for 20 mins before collecting total RNA for RT-qPCR.

Sequence of dsRNA used in the study are:

dsGFP_Fwd: TAATACGACTCACTATAGGGGAGGAGCTGTTCACC

dsGFP_Rev: TAATACGACTCACTATAGGGGGCGAGCTGCACGCTGCC

Mal-d-dsRNA-1_Fwd: TAATACGACTCACTATAGGGCAAGTCAGCGATGTTCTGGA

Mal-d-dsRNA-1_Rev: TAATACGACTCACTATAGGGTTTGGCTTTTCACTACCGCT

Mal-d-dsRNA-2_Fwd: TAATACGACTCACTATAGGGGCAAGTCCGGTCAATCATCT

Mal-d-dsRNA-2_Rev: TAATACGACTCACTATAGGGTTTTGCTGCTCCAACATCAG

Mal-d-dsRNA-3_Fwd: TAATACGACTCACTATAGGGGCAGCAACTTGAGATGGACA

Mal-d-dsRNA-3_Rev: TAATACGACTCACTATAGGGAAAGTGACTATTCACCGCCG

Mal-d-dsRNA-4_Fwd: TAATACGACTCACTATAGGGTGCCAAGCTCAATCACTTTG

Mal-d-dsRNA-4_Rev: TAATACGACTCACTATAGGGAGGACTGTTGCCATTTCCAG

### Real-time quantitative PCR

#### in fly ovaries

For heat shock experiments, seven pairs of ovaries per genotype were dissected from five days old female flies in Schneider’s drosophila media and placed at a +37°C waterbath for 20 mins. Ovaries were snap frozen in liquid Nitrogen and homogenised to extract total RNA with Nucleospin RNA II kit.

#### in S2R+ cells

For heat shock experiments in S2R+ cells, drugs were added to the Schneider’s full media for defined timepoints (16 h and 10 min) at the following concentration: DMSO (0.25 %), CCG-1423 (10 uM) and CG203971 (25 uM). Then, heat shock for S2R+ cells was done by placing sealed culture flasks into a + 37°C water bath for 20 mins or cells were incubated at +25°C for the same time. After that cells were immediately lysed to extract total RNA with Nucleospin RNA II kit.

For ecdysone stimulation S2R+ cells were treated with 1.5 uM of 20-E in Schneider’s full media for 1 h following 10 min drug treatment at the concentration: DMSO (0.1 %), CCG-1423 (10 uM) and ActD (5 uM). After that cells were immediately lysed to extract total RNA with Nucleospin RNA II kit.

#### in NIH 3T3 cells

For heat shock experiments in NIH 3T3 cells culture dishes were placed in + 42°C water bath for 20 mins, following concentrations of the drugs were added to DMEM media for defined timepoints (16 h or 10 min): DMSO (0.4 %), CCG-1423 (10 uM), CCG-203971 (40 uM).

Pirin inhibition was performed by treating NIH 3T3 cells with pirin inhibitor CCT251236 for 10 mins, followed by heat shock at + 42°C water bath for 20 mins or serum stimulation (15% serum) for 20 min. The concentration of inhibitors used for heat shock experiment were 5 nM, 10 nM and 100 nM and DMSO (0.4%). The concentration of inhibitors used for serum stimulation experiments were 5 nM, 10 nM, 20 nM, 50 nM or 100 nM and DMSO (0.4 %).

For serum stimulation in NIH 3T3 (MRTF_KO-1 and MRTF_KO-2) and wild type NIH 3T3 (MRTF_WT) was performed by culturing cells overnight in DMEM containing 0.3 % of FBS. After starvation, cells were stimulated with serum (15 %) for 45 min and harvested for RNA extraction.

#### RNA extraction and qPCR

Total RNA was extracted and purified using NucleoSpin RNA kit according to the manufacturer’s protocol; RNA purification from cultured cells and tissue (740955; Macherey-Nagel). Complementary cDNA was synthesised using Maxima first strand cDNA kit from five hundred nanograms of purified total RNA (K1641; ThermoFisher Scientific). SYBR green dye based (BIO-98005; Meridian Bioscience) qPCR of target genes were performed in the Bio-Rad CFX machine (Bio-Rad). Relative expression levels were calculated by the comparative *C*_T_ method, normalising to the Gapdh cDNA: 2^−C^_T_ (target)/2^−C^_T_ (Gapdh) for mammalian cells and Rpl32 cDNA: 2^−C^_T_ (target)/2^−C^_T_ (Rpl32) for fly cells and ovaries.

**Supplementary table 2.**
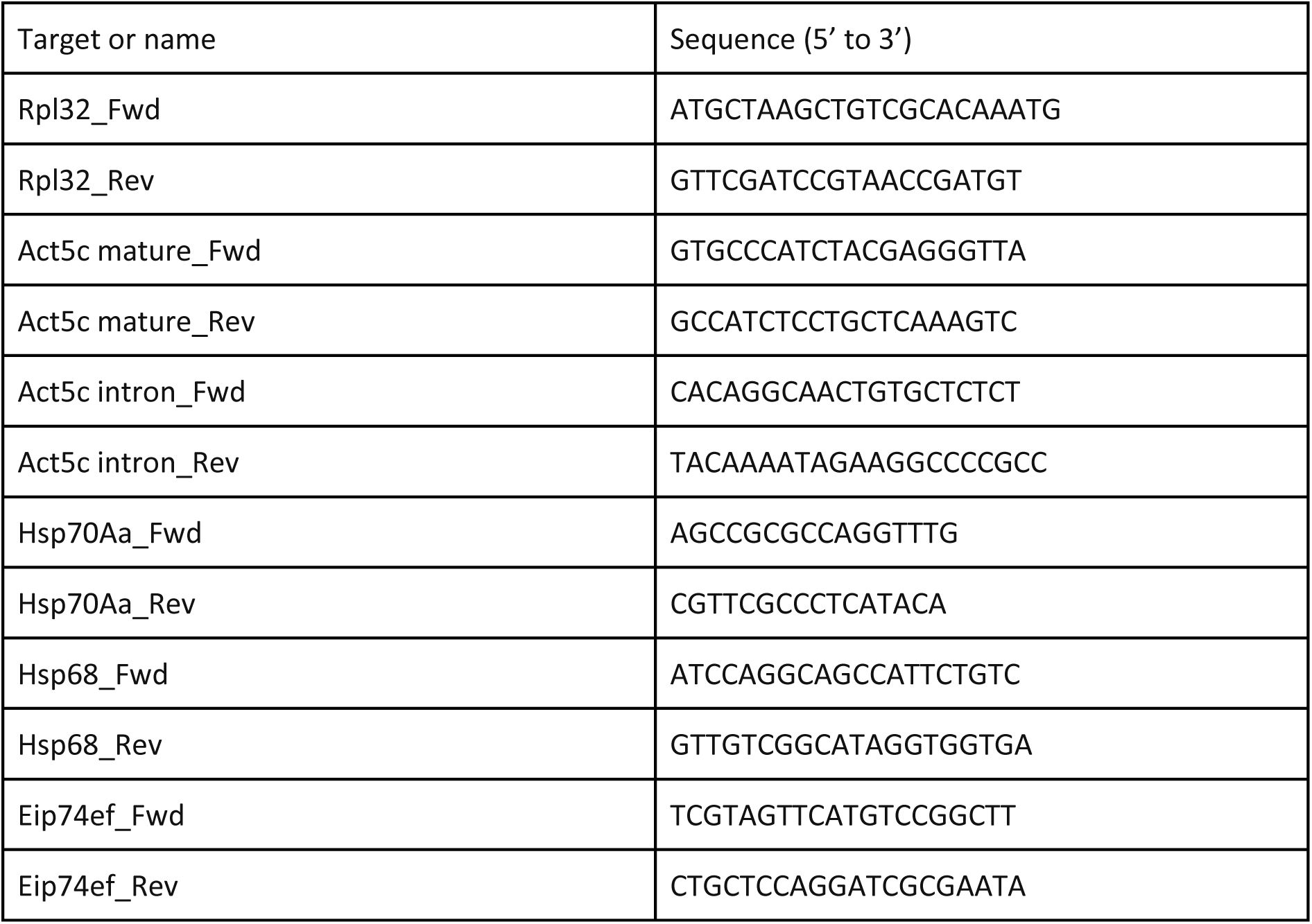
(Primers for qPCR in Drosophila melanogaster ovaries and S2R+ cells)

**Supplementary table 3.**
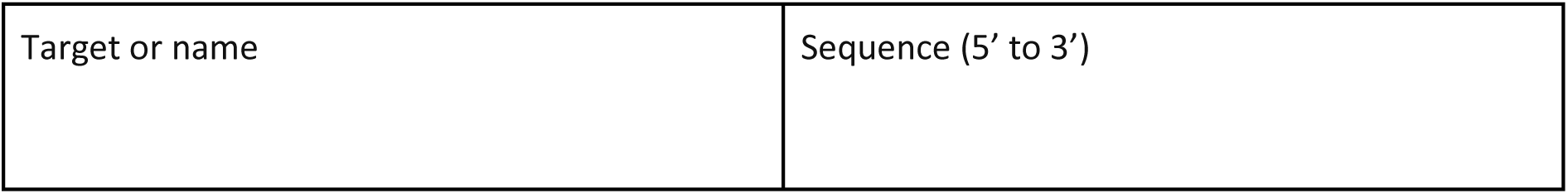
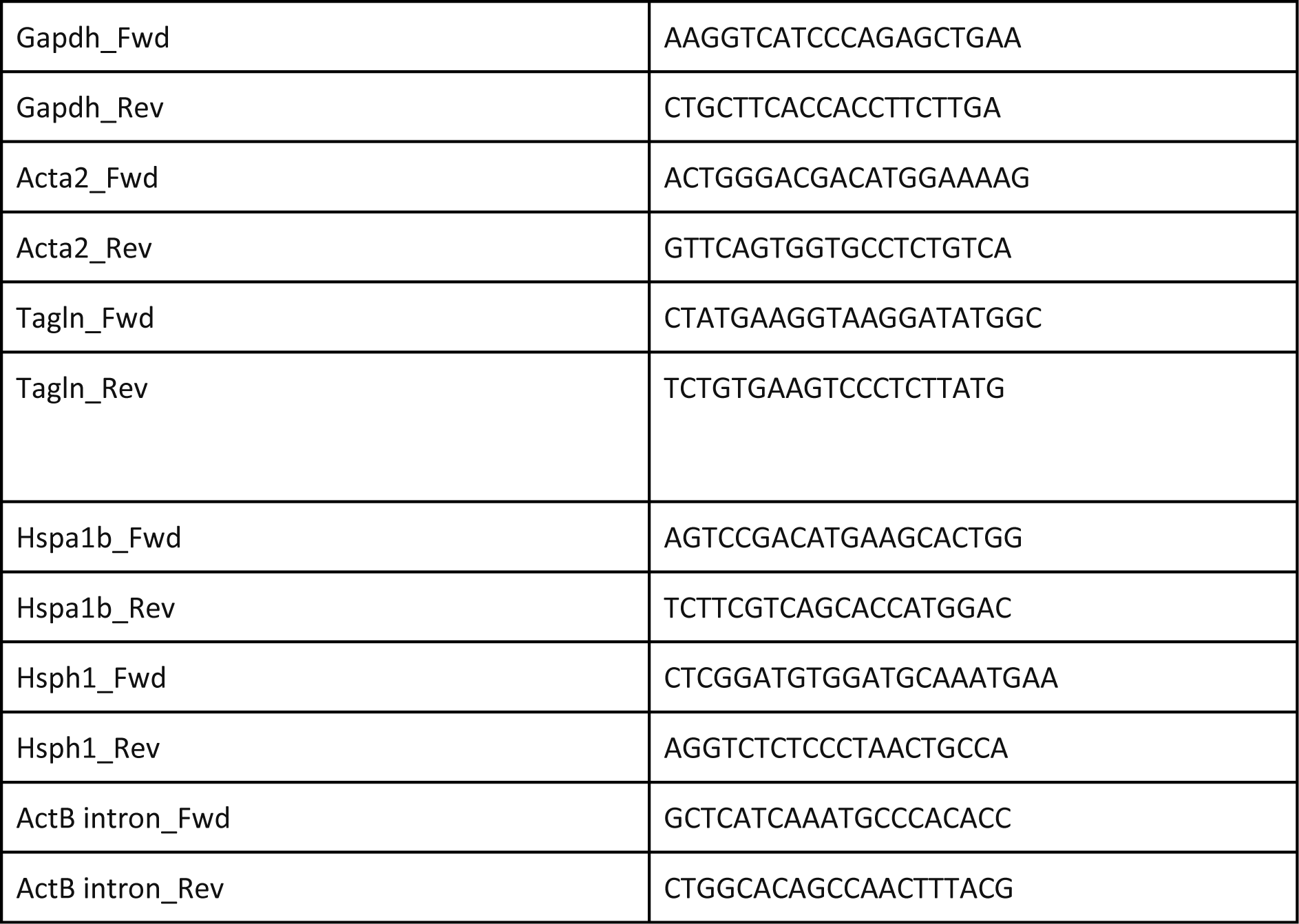
(Primers for qPCR in NIH3T3 cells)

### Luciferase assay

Luciferase assay was done in a 24 wells plate format. Overnight cultured NIH3T3 cells were transfected with 8 ng of SRF reporter plasmid p3DA.luc, 20 ng of reference reporter pRL-TK and pEF-Myc to a final amount of 200 ng by using JetPrime transfection reagent. After 7 h of transfection, DMEM was changed to 0.3 % FBS and treated with CCG-1423 (10 uM), CCG-100602 (40 uM) and CCG-203971 (40 uM). After 16 h, serum stimulation was done with DMEM containing 15 % FBS and cells were harvested after 7 h of stimulation and SRF reporter activity was measured with Dual-Luciferase reporter assay system (E1910; Promega) and a luminometer, according to manufacturer’s instructions. For data analysis, the activity of firefly luciferase was normalised to the renilla luciferase activity.

### 5-EU labelling, staining and microscopy

Click-iT™ RNA Alexa Fluor™ 488 Imaging was purchased from Invitrogen (C10329). Additional 5-Ethynyl-uridine (5-EU) was purchased from Jena Bioscience (CLK-N002) and resuspended to an initial stock concentration of 400 mM in DMSO.

For EU labelling into cells, cells were grown on glass coverslips (13 mm diameter, Thickness No. 1.5) (631-0150; VWR) overnight. For S2R+ cells DMSO (0.25 %), CCG-1423 (10 uM), CG203971 (25 uM), ActD (5 uM) and for NIH 3T3 cells DMSO (0.4 %), CCG-1423 (10 uM), CCG-203971 (40 uM), ActD (5 uM) were added for 10 minutes. After inhibitor treatment, 5-EU (2 mM) was added to cells for 20 mins (S2R+) or 30 mins (NIH3T3). Cells were fixed with 4 % paraformaldehyde (PFA), permeabilized and then proceeded for EU detection with Click chemistry using A488-azide according to the staining protocol from the manufacturer.

EU labelled cells were co-stained with anti-Histone H4 pAb (1:250 in Dulbecco wash buffer with 0.2 % w/v BSA) (SAB4500311; Sigma Aldrich) for 30 mins at room temperature in a dark humidified chamber. Cells were rinsed in Dulbecco wash buffer, stained with secondary antibody for 20 mins, and finally mounted on glass slide with ProLong Diamond antifade reagent. Secondary antibody used in this study is Alexa-Fluor-647-conjugated anti-rabbit pAB (A32733; Invitrogen).

All fluorescence images were collected by Leica Stellaris 8 Falcon confocal microscope using the Leica Application Suite (LAS X) software (version 4.3.0). The HC PL APO CS2 40x/1.25 GLYC objective and White laser light (WLL) was used to image the samples. Detection of the fluorescence was done with HyD S for signals of 5-EU, excited at 488 nm and HYD X for the signals of Histone H4 excited at 649 nm. The voxel size for each individual image was 0.1893×0.1893×0.3600 micron^3 and pinhole 1 AU was selected.

Images of the nucleus were obtained by randomly selecting fields as assessed by histone H4 staining. Z-stacks of the individual samples were collected with the same settings for control (DMSO) and treated (CCG-1423, CCG-203971 and Actinomycin-D treatment) cells. Maximum projections of confocal planes through the entire image were used and areas of interest were drawn based on histone H4 staining in the nucleus, excluding the bright nucleolar dots. After this, mean intensities of 5-EU labelled nascent RNA were quantified within this area. All image analyses were carried out using ImageJ software.

### ChIP seq

Sample preparation for ChIP-seq was based on previous protocols (Sidorenko et al., 2022; Sokolova et al., 2018). In short, cells were plated in T75 flasks (S2R+) or 15 cm plates (NIH3T3). The next day, drug treatment was done for allocated timepoints (10 mins with the inhibitors at NH condition) followed by heat shock for 20 mins either in 37°C waterbath (S2R+ cells) or 42°C waterbath (NIH 3T3 cells). Cells were crosslinked with formaldehyde for 10 mins after drug/heat shock treatments. Crosslinking was quenched with ice-cold 125 mM glycine on ice for five mins. Chromatin fragments were obtained from RIPA buffer lysed cells by sonication with Diagenode Bioruptor (15 cycles, 30 sec on and 30 sec off, highest power setting). Immunoprecipitation was carried out with 5 ug of antibody overnight at 4°C. The immune-complexes were collected with 50 ul of protein A sepharose (17-0780-01; GE Healthcare) at 4°C for two hours with rotation. Chromatin was eluted from washed beads in 1% SDS in TE buffer (10 mM Tris-HCl, pH 8.0), 1 mM EDTA). DNA purification after reverse crosslinking was done with phenol/chloroform/isoamyl alcohol (17908; ThermoFisher Scientific) and resuspended in 80 ul of nuclease free water.

ChIP libraries were prepared for Illumina NextSeq 500 using NEBNext ChIP-Seq DNA Sample Prep Master Mix Set for Illumina (E6240; NEB) and NEBNext® Multiplex Oligos for Illumina® (Index Primers Set 1) (E7335; NEB) according to the manufacturer’s protocols. Sequencing was performed with NextSeq500 at Biomedicum Functional Genomics Unit (FuGU). ChIP-Seq data sets were aligned using Bowtie2 (using Chipster software)(Kallio et al., 2011) to version mm10 of the mouse genome and dm6 of Drosophila genome with the default settings. To analyse, visualise and present ChIPseq data, we used Integrative Genomics Viewer (IGV) (Robinson et al., 2011), deepTools (Ramirez et al., 2016), public server at usegalaxy.org (Galaxy, 2022) and EaSeq (http://easeq.net) (Lerdrup et al., 2016). The signals from the regions of interest were quantified with normalization per million per kilobase (RPKM). Signals from the gene bodies were normalised to the size of the region. For Drosophila TSS regions were defined from −250bp to +250bp, CPS regions are from 0 to +1000bp and gene bodies as +250bp from TSS till CPS based on the dm6 version. For mouse TSS regions were defined from −100bp to +400bp, CPS regions are from 0 to +4000bp and gene bodies as +400bp from TSS till CPS based on the mm10 version. Statistical analysis was performed with Mann-Whitney U test using Origin from OriginLab.

### PRO-seq library generation

Sample preparation for PRO-seq was conducted as previously described with slight modifications (Chu et al., 2018; Mahat et al., 2016a; Vihervaara et al., 2021). Chromatin from 10 million S2R+ cells was isolated with NUN buffer (0.3 M NaCl, 1 M Urea, 1 % Igepal, 20 mM HEPES pH 7.5, 7.5 mM MgCl2, 0.2 mM EDTA, 1 mM DTT, 20 units/ml Superase In RNase Inhibitor (A2696; Invitrogen) and 1 x Protease Inhibitor (4693132001; Sigma Aldrich)), and fragmented with Diagenode Bioruptor using the high throughput power setting, 30 s ON/ 30 s OFF, for five cycles. The solubilized chromatin was flash-frozen and stored at −80°C. Before run-on reaction, an equal amount of solubilized chromatin from untreated NIH3T3 cells was spiked into each sample, (counted to account for 1 % of the total chromatin). The run-on reaction was conducted at 37°C for 5 min in the presence of biotinylated nucleotides (10 mM Tris-HCl pH 8.0, 5 mM MgCl2, 300 mM KCl, 1 mM DTT, 0.04 mM biotin-A/C/G/UTP (R0481; ThermoFisher Scientific), 20 U/μl SUPERase In RNase inhibitor and 1 % Sarkosyl). Trizol LS (10296010, ThermoFisher Scientific) was used to extract total RNA and EtOH was used to precipitate the RNA. The RNA was base hydrolyzed with 1 N NaOH for 5 min on ice, and NaOH neutralised with Tris-HCl (pH 6.8). Unincorporated biotin-NTPs were removed with RNase free P-30 columns (7326223; Bio-Rad), and three cycles of magnetic MyOne Streptavidin C1 bead purification (65002; Invitrogen) was performed to isolate the biotinylated nascent transcripts. After the first bead purification, nascent RNA was isolated with streptavidin beads and barcoded adapters ligated at 3’ ends of the nascent RNA (T4 RNA ligase 1 enzyme, M0204L; NEB) overnight at 25° C. After the second streptavidin bead-binding, 5’-decapping (RppH, M0356S; NEB) and 5′-hydroxyl group repair (T4 polynucleotide kinase (M0201L; NEB) were conducted on beads. Transcripts were purified with Trizon and EtOH, and sfter the third bead-binding, reverse transcribed with Superscript III Reverse Transcriptase enzyme (18080-044; Invitrogen). Sequencing libraries were generated using Phusion DNA polymerase with TruSeq small-RNA adaptors and sequenced at NOVOGene USA using NextSeq500 (Illumina).

### Computational analyses of PRO-seq data

The PRO-seq reads were adaptor-clipped using cutadapt (Martin, 2011)and trimmed and filtered with fastx toolkit (http://hannonlab.cshl.edu/fastx_toolkit/). Reads were aligned to the *Drosophila* (dm6) and mouse (mm10) genomes using Bowtie 2 (Langmead and Salzberg, 2012). Reads that uniquely mapped to the *Drosophila* (dm6) genome, were retained. The reads that uniquely mapped to the mm10 chromosomes provided a count of reads for spike-in derived normalization factors.

Mapped reads were processed from bed files to coverage files, retaining only the 3′ end nucleotide (active sites of transcription) of each read. Density normalized bedgraph files were adjusted by sample-specific normalization factors as described previously (Booth et al., 2018; Himanen et al., 2022).

For PRO-seq analyses, gene regions were defined as previously described (Rabenius et al., 2022): promoter-proximal region as −250nt to +250nt from TSS; gene body as +251nt from the TSS to −500nt from the CPS, and termination window, due to gene-dense *Drosophila* genome, as 0 to +1000nt from the CPS. The level of transcription *per* each annotated transcript was measured from the gene body, using spike-in derived size factors (Booth et al., 2018; Vihervaara et al., 2021) with DESeq2 (Love et al., 2014). The analyses of enriched gene annotation categories were performed with Database for Annotation, Visualization and Integrated Discovery (DAVID; (Dennis et al., 2003)).

### Statistical analysis

Statistical analyses were performed using Origin PRO version 2021b sr2. Statistical significance was determined by two-tailed Student’s *t*-test, with two-sample equal variance for the data conformed to normal distribution (qPCR analysis of heat shock inducible genes, MRTF/SRF target genes and ecdysone inducible genes). For 5 Ethynyl uridine stained imaging data, distribution of the data (mean grey value) was checked using Shapiro-Wilks test in Origin PRO version 2021b sr2. Since the distribution didn’t follow the normal distribution curve, data was analysed using a non-parametric test (Mann-Whitney U test) at significance level 0.05 with P<0.05 rejecting the null hypothesis. P value >0.05 represents non significance.

All graphs in the figure panels in the manuscript represent three or more independent biological repetitions. PRO-seq data were repeated in two independent experiments/biological replicates.

## Supplementary figures

**Figure S1:**
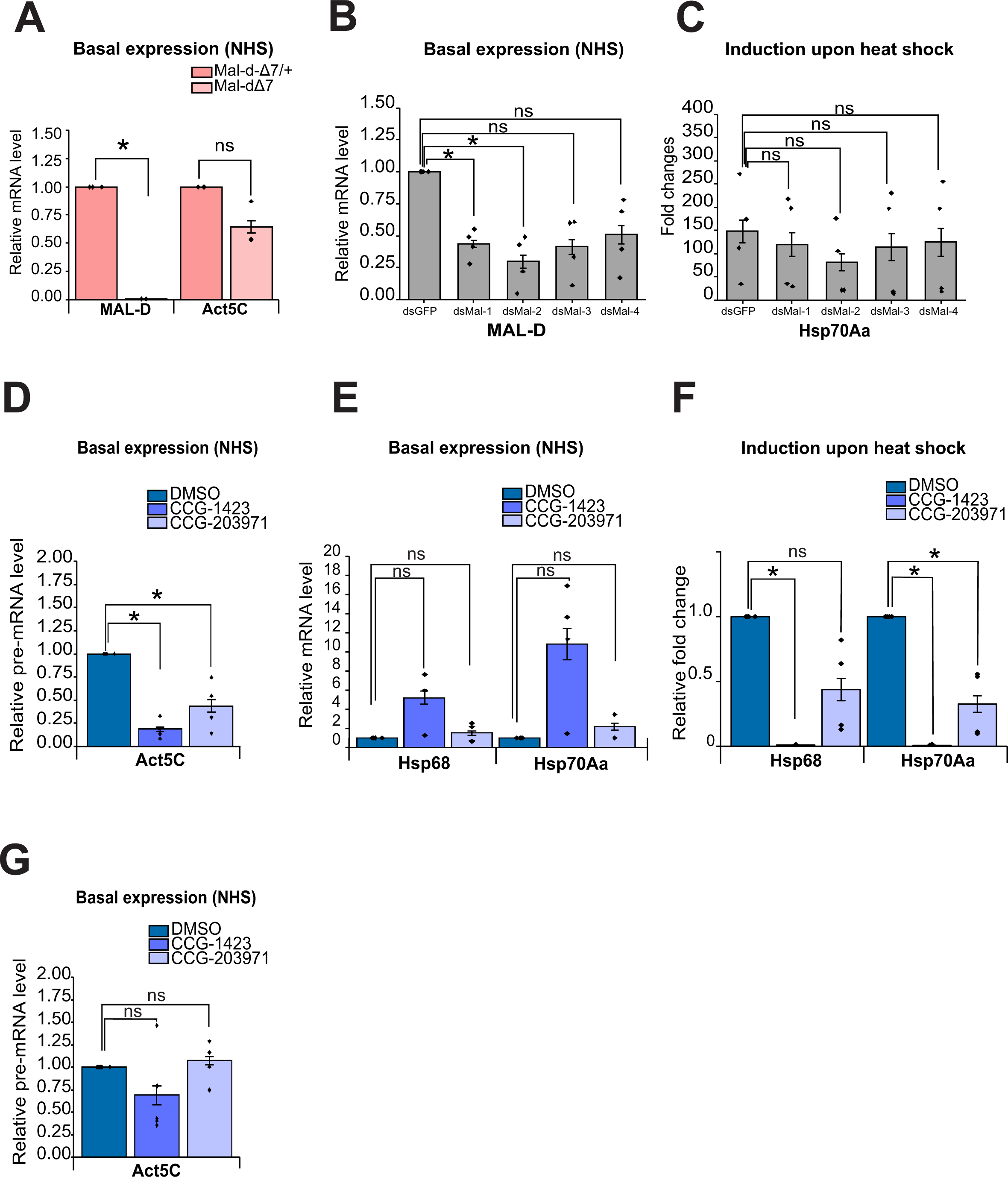
Deletion of MAL-D and CCG-1423-derived inhibitors reduce *hsp* gene induction upon HS. Related to Figure 1. **A.** Validation of MAL mutant flies. Expression of *MAL-D* and *Act5C* mRNAs in heterozygous (mal-dΔ7/+) and homozygous (mal-dΔ7) MAL-D mutant fly ovaries. Data is from three biological replicates and normalized to mal-dΔ7/+ values. The bar indicates the mean, individual data points are shown and error bars are standard errors of the mean (s.e.m). Statistical significances with student’s one-way t-test (*; P< 0.05). P-values: Mal-dΔ7/+ vs Mal-dΔ7 for Mal-d is 8E-7, Mal-dΔ7/+ vs Mal-dΔ7 for Act5c is 0.09. **B.** Knockdown efficiency of double stranded RNAs (dsRNAs) targeting different regions of MAL-D. Expression levels of *MAL-D* mRNAs in S2R+ fly cells treated with non-targeting dsRNA (dsGFP) and dsRNAs targeting MAL-D (dsMal-1-4). Data is from four biological replicates and normalized to dsGFP values, and shown as in A. Statistical significance with student’s one-sample t-test (*; P< 0.0125 with Bonferroni correction). P-values: dsMal-1 vs dsGFP is 0.002, dsMal-2 vs dsGFP is 0.002, dsMal-3 vs dsGFP is 0.016 and dsMal-4 vs sGFP is 0.039. **C.** Effect of MAL-D dsRNAs on *Hsp70Aa* mRNA induction. Fold changes in *Hsp70Aa* induction upon heat shock in S2R+ fly cells treated with MAL-D dsRNAs as in B. Data is from four biological replicates,and shown as in A. Statistical significance with one-way ANOVA followed by Tukey’s multiple comparison test (*; P< 0.0125 with Bonferroni correction). P-values: dsMal-1 vs dsGFP is 0.99, dsMal-2 vs dsGFP is 0.89, dsMal-3 vs dsGFP is 0.99 and dsMal-4 vs dsGFP is 0.99. **D.** Effect of 16h incubation with CCG-derived compounds on *Act5C* expression. Expression levels of *Act5C* mRNA in S2R+ cells treated with DMSO, CCG-1423 and CCG-203971 for 16 h. Data is from four biological replicates, normalized to DMSO values and shown as in A. Statistical significance with one-sample student’s t-test (*; P< 0.025 with Bonferroni correction). P-values: CCG-1423 vs DMSO is 0.006 and CCG-203971 vs DMSO is 0.024. **E.** Effect of 16h incubation with CCG-derived compounds on baseline expression of *hsp* genes. Expression level of *Hsp68* and *Hsp70Aa* mRNAs in S2R+ cells treated with DMSO, CCG-1423 and CCG-203971 for 16 h. Data is from four biological replicates, normalized to the DMSO values and shows as in A. Statistical significance with student’s one-sample t-test (*; P< 0.025 with Bonferroni correction). P-values: CCG-1423 vs DMSO for *Hsp68* is 0.055, CCG-203971 vs DMSO for *Hsp68* is 0.39, CCG-1423 vs DMSO for *Hsp70Aa* is 0.06 and CCG-203971 vs DMSO for *Hsp70Aa* is 0.19. **F.** Effect of 16h incubation with CCG-derived compounds on heat shock induction of *hsp* genes. Fold change in *Hsp68* and *Hsp70Aa* mRNAs upon 30 min heat shock in S2R+ cells treated with DMSO, CCG-1423 and CCG-203971 for 16 h. Data is from four biological replicates and shown as in A. Statistical significance with student’s one-sample t-test (*; P< 0.025 with Bonferroni correction). P-values: CCG-1423 vs DMSO for *Hsp68* is 3E-8, CCG-203971 vs DMSO for *Hsp68* is 0.047, CCG-1423 vs DMSO for *Hsp70Aa* is 8E-8 and CCG-203971 vs DMSO is 0.013. **G.** Effect of 30 min incubation with CCG-derived compounds on *Act5C* expression. Expression level of *Act5C* in S2R+ cells treated with DMSO, CCG-1423 and CCG-203971 for 30 min. Data is from five biological replicates, normalized to DMSO values and shown as in A. Statistical significance with student’s one-sample t-test (*; P< 0.025 with Bonferroni correction). P-values: CCG-1423 vs DMSO is 0.212, CCG-203971 vs DMSO is 0.468.

**Figure S2.**
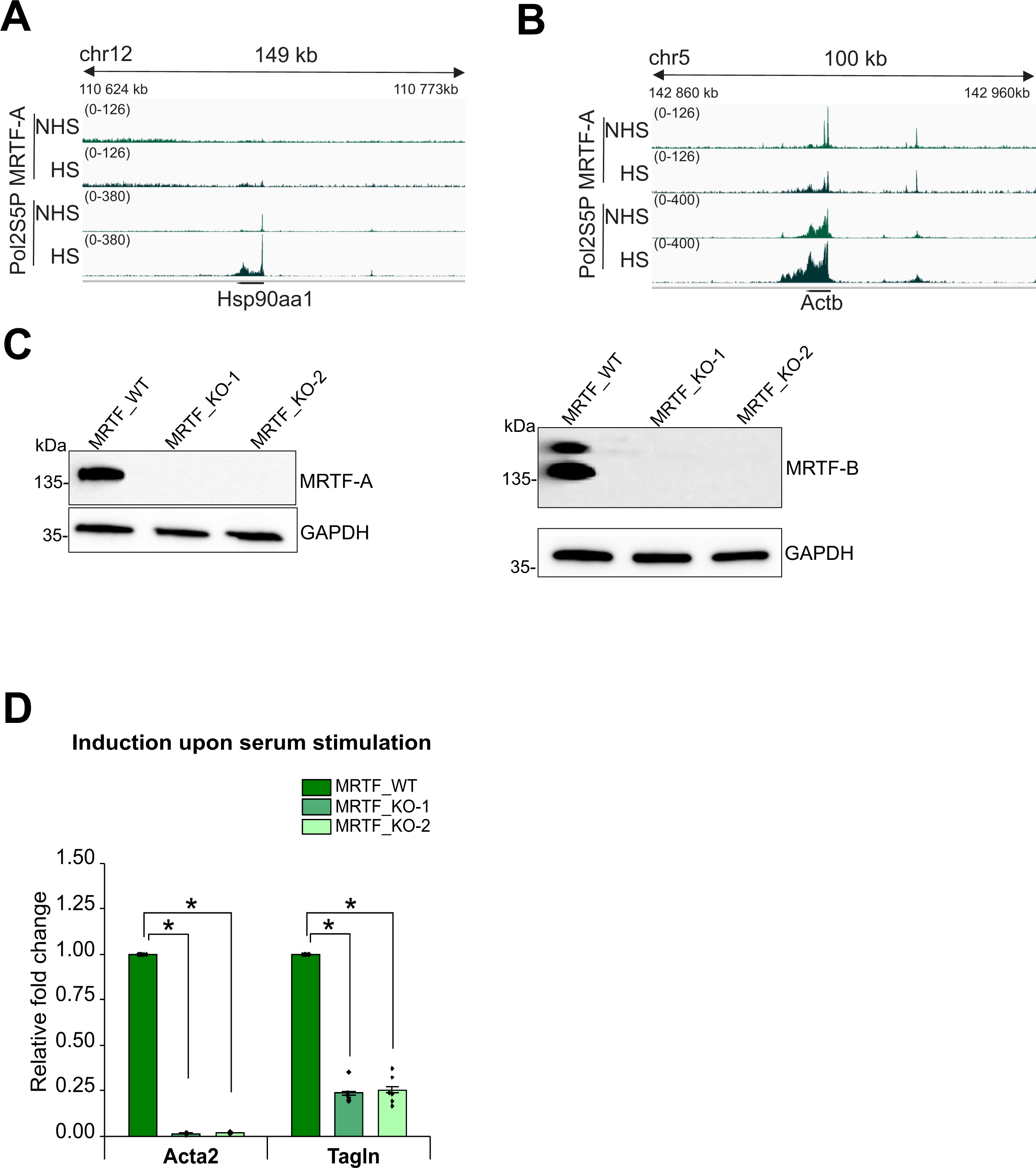
Analysis of Hsp gene transcriptional regulation by MRTF in mouse fibroblasts. Related to figure 2. **A.** Normalized (Reads Per Kilobase per Million mapped reads (RPKM)) coverage of MRTF-A and RNA polymerase II phosphorylated on serine 5 (Pol2S5P) on heat shock responding gene *Hsp90aa1* in mouse NIH 3T3 cells in non-heat shock (NHS) and heat shock (HS) conditions from ChIP-seq. **B.** Normalized (RPKM) coverage of Pol II S5P and MRTF-A on MRTF/SRF target gene *Actb* in NIH 3T3 cells in non heat shock (NHS) and heat shock (HS) conditions. **C.** Western blot showing the absence of MRTF-A and MRTF-B proteins in MRTF-KO clones. Name of the clones are on the top, molecular weight markers on the left, and antibodies used in Western Blot on the right. GAPDH was used as a loading control. **D.** Lack of MRTF in NIH 3T3 cells significantly reduces the induction of serum responsive genes. Fold changes in *Acta2* and *Tagln* mRNA induction upon serum stimulation in MRTF deleted (MRTF_KO-1 and MRTF_KO-1) and MRTF wildtype (MRTF_WT) NIH 3T3 cells. Data is from seven (*Acta2*) or six (*Tagln*) biological replicates and normalized to MRTF_WT values. Bars represent the mean, individual data points are shown and the error bars represent s.e.m. Statistical significance with Student’s one-sample t-test (*; P< 0.025 with Bonferroni correction). P-values for *Acta2*: MRTF_WT vs MRTF_KO-1 is 1E-14, MRTF_WT vs MRTF_KO-2 is 1E-10. P-values for *Tagln*: MRTF_WT vs MRTF_KO-1 is 6E-7, MRTF_WT vs MRTF_KO-2 is 3E-6.

**Figure S3.**
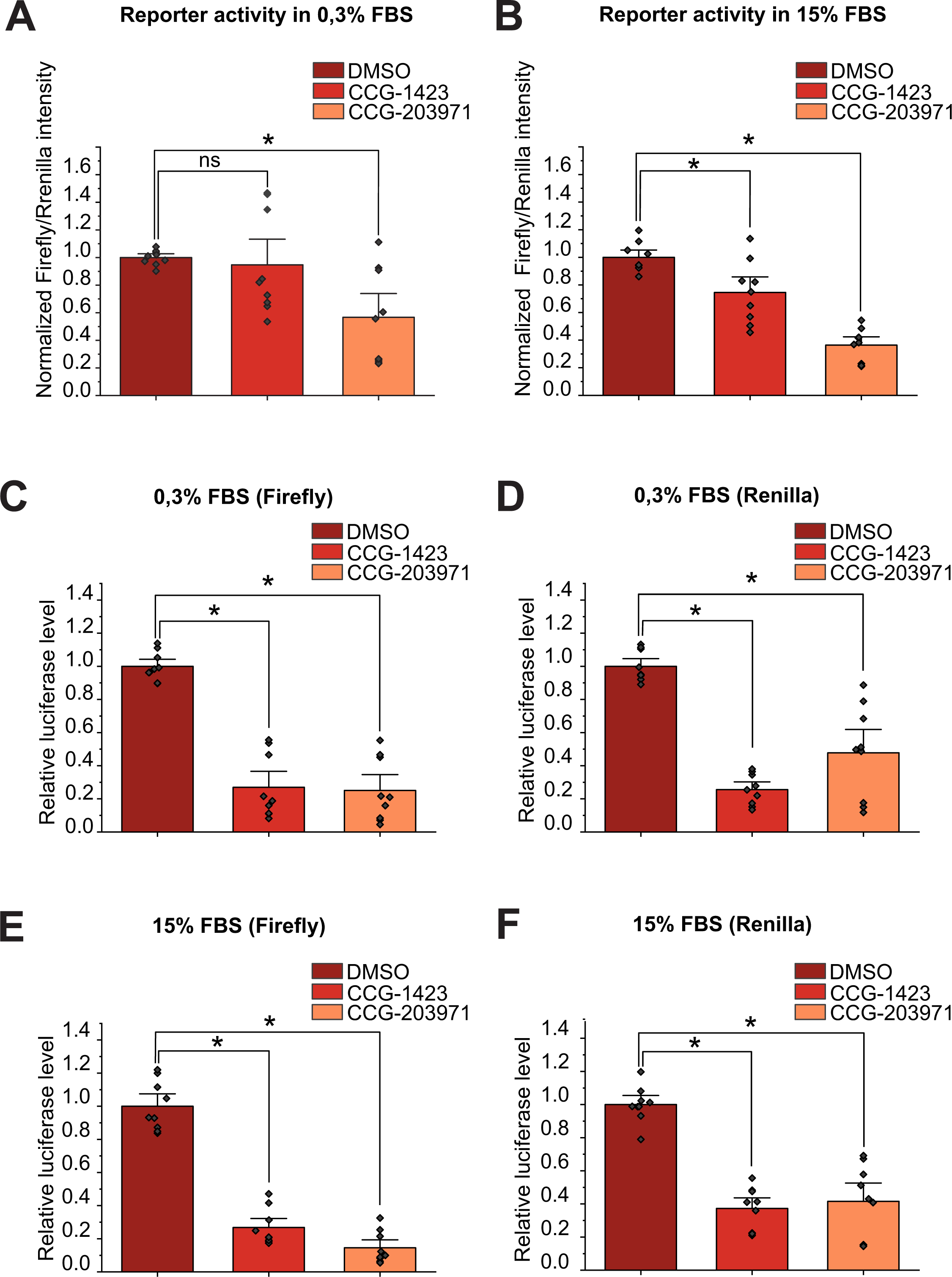
The effect of CCG-derived inhibitors is not limited to SRF-reporter genes. Related to Figure 3. **A.** SRF reporter assay with serum response element (SRE)-controlled firefly luciferase activity normalized to constitutive renilla luciferase activity in cells treated with DMSO, CCG-1423 and CCG-203971 in serum starved conditions. Data is from three biological replicates, bars represent the mean, normalized to the mean of the DMSO values, individual data points indicated and error bars are s.e.m. Statistical significance with one-way ANOVA followed by Tukey’s multiple comparison test (*; P< 0.05). P-value: CCG-1423 vs DMSO is 0,92, CCG-203971 vs DMSO is 0.012. **B.** Normalized SRF reporter activity in serum-stimulated conditions shown as in A. Statistical significance with one-way ANOVA followed by Tukey’s multiple comparison test (*; P< 0.05). P-values: CCG-1423 vs DMSO is 0.007, CCG-203971 vs DMSO is 1E-00. **C.** SRE-controlled firefly luciferase activities in serum starved condition shown as in A. Statistical significance with one-way ANOVA followed by Tukey’s multiple comparison test (*; P< 0.05). P-values: CCG-1423 vs DMSO is 1E-00, CCG-203971 vs DMSO is 1E-00. **D.** SRE-controlled firefly luciferase activities in serum stimulated conditions shown as in A. Statistical significance with one-way ANOVA followed by Tukey’s multiple comparison test (*; P< 0.05).P-values: CCG-1423 vs DMSO is 1E-00, CCG-203971 vs DMSO is 6E-6. **E.** Renilla luciferase activities in serum starved condition shown as in A. Statistical significance with one-way ANOVA followed by Tukey’s multiple comparison test (*; P< 0.05). P-values: CCG-1423 vs DMSO is 1E-00, CCG-203971 vs DMSO is 1E-00. **F.** Renilla luciferase activities in serum stimulated condition shown as in A. Statistical significance with one-way ANOVA followed by Tukey’s multiple comparison test (*; P< 0.05). P-values: CCG-1423 vs DMSO is 1E-00, CCG-203971 vs DMSO is 1E-7.

**Figure S4.**
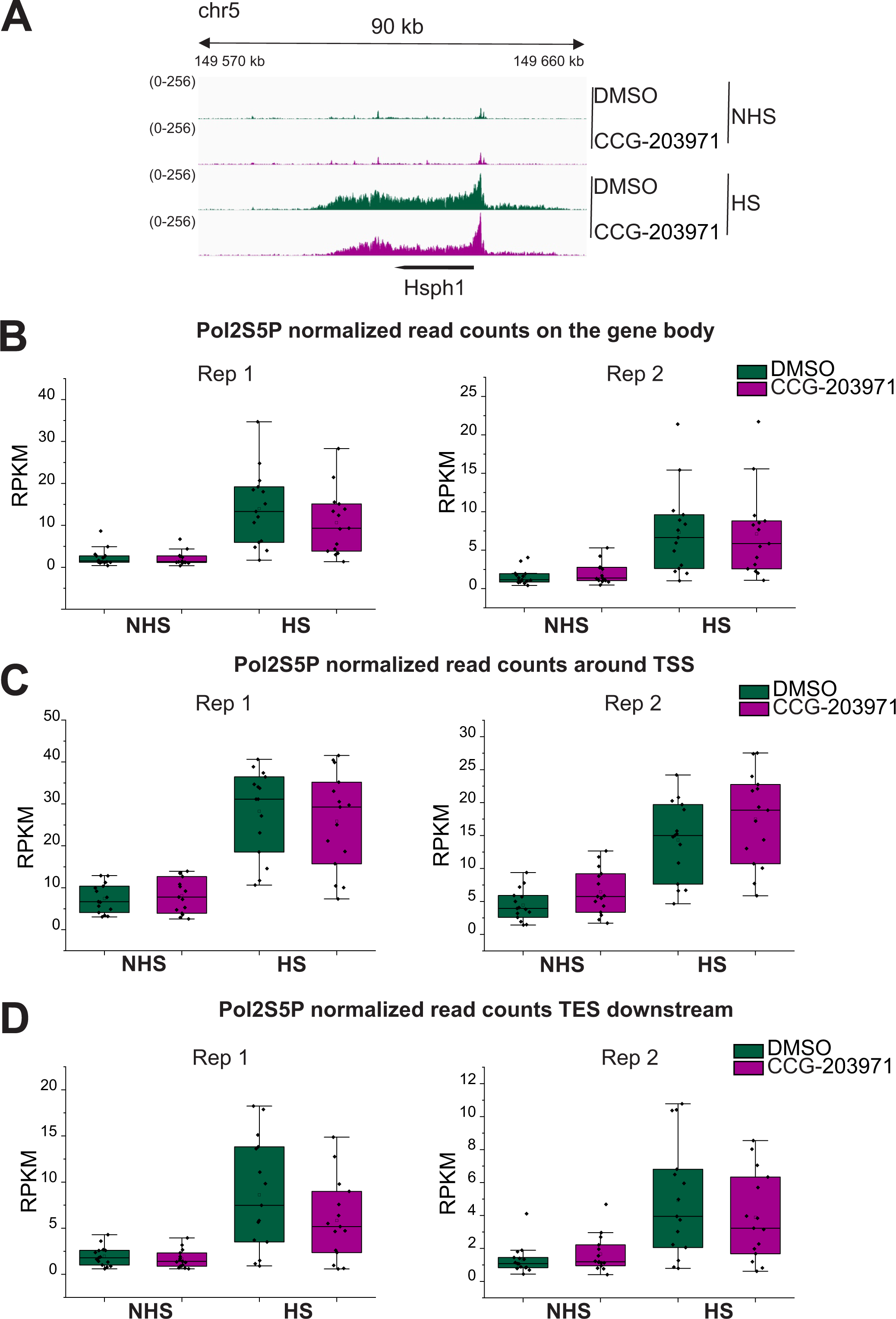
Binding of RNA polymerase II to *hsp* genes in CCG-203971 treated mouse fibroblasts. Related to figure 4. A. Normalized (RPKM) coverage of RNA polymerase II phosphorylated on serine 5 (Pol2S5P) on heat shock responding gene *Hsph1* in NIH 3T3 mouse cells in DMSO and CCG-203971 treated cells in non-heat shock (NHS) and heat shock (HS) conditions from ChIP-seq. B. Quantification of Pol2S5P on the gene body of the 15 *hsp* genes as normalized read counts (RPKM) in DMSO and CCG-203971 treated cells in non-heat shock (NHS) and heat shocked (HS) conditions in two replicate experiments. Data is shown as a 25th and 75th percentiles box plot, where the horizontal line represents the median, small box represents mean and whiskers represent data range within 1.5IQR. Statistical significance with Mann-Whitney U test (*; P< 0.05). C. Quantification of Pol2S5P around the transcription start site (TSS) of the *hsp* genes shown as in B. D. Quantification of Pol2S5P downstream of the CPS of the *hsp* genes shown as in B.

**Figure S5.**
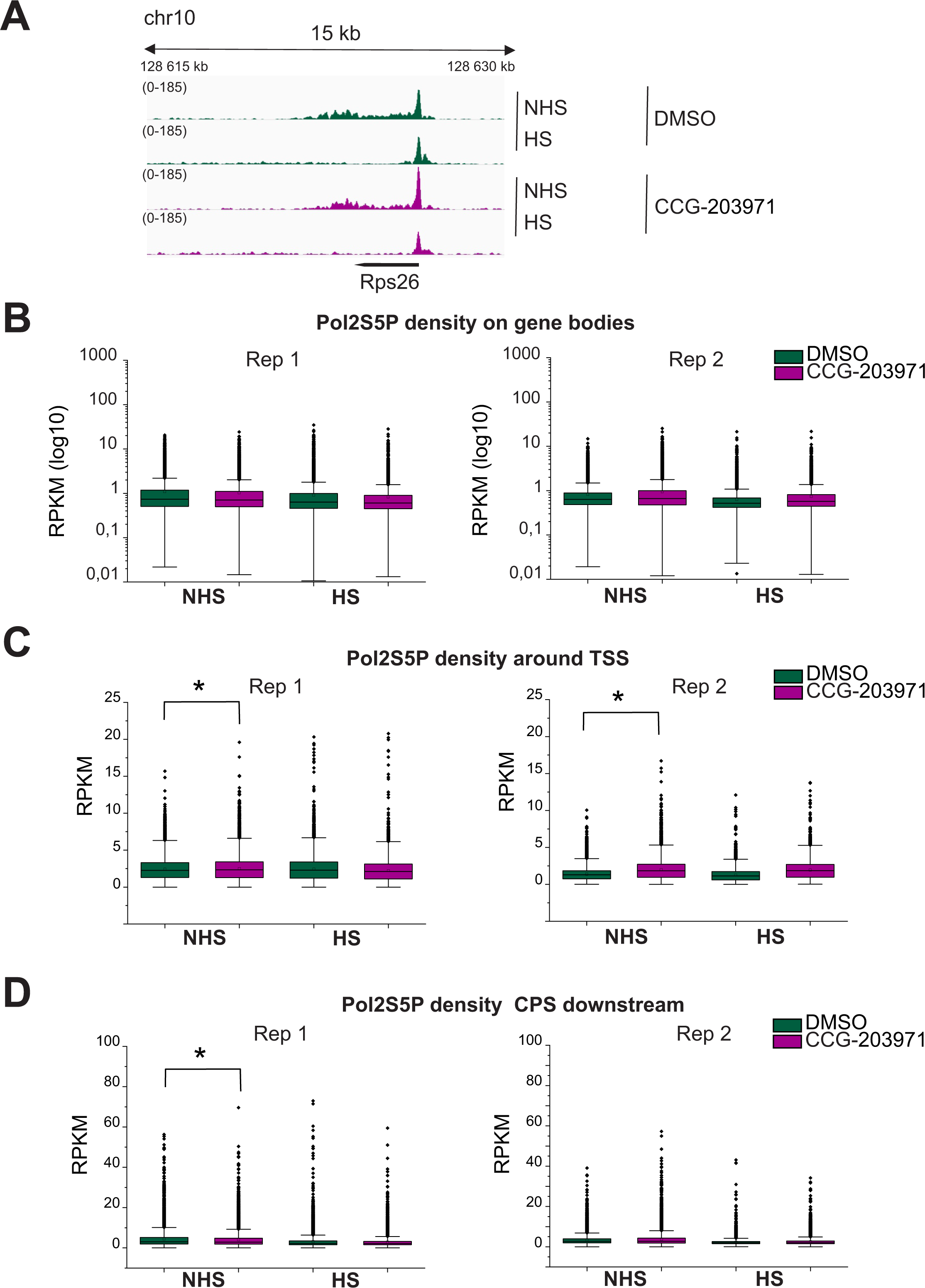
CCG-203971 results in accumulation of RNA polymerase II on the transcription start site in mouse fibroblasts **A.** Normalized (RPKM) coverage of RNA polymerase II phosphorylated on serine 5 (Pol2S5P) on the *Rps26* gene in NIH 3T3 mouse cells in DMSO and CCG-203971 treated cells in non-heat shock (NHS) and heat shock (HS) conditions from ChIP-seq. **B.** Quantification of Pol2S5P on the gene body of the 11468 expressed genes in NIH3T3 mouse cells in DMSO and CCG-203971 treated cells in non-heat shock (NHS) and heat shock (HS) conditions from two replicate experiments. Data is shown as a 25th and 75th percentiles box plot, where the horizontal line represents the median, small box represents mean and whiskers represent data range within 1.5IQR. Statistical significance was calculated by Mann-Whitney U test. **C.** Quantification of Pol2S5P around the transcription start site (TSS) of the expressed genes shown as in B. Statistical significance was calculated by Mann-Whitney U test. (*; P< 0.05). P-values for NHS DMSO vs CCG-1423 in replicate 1 is 0.0183 and in replicate 2 is 3,9E-256. **D.** Quantification of Pol2S5P downstream of transcription end site (CPS) of the expressed genes shown as in B. Statistical significance was calculated by Mann-Whitney U test. (*; P< 0.05). P-values for NHS DMSO vs CCG-1423: replicate 1 is 0.000001.

**Figure S6.**
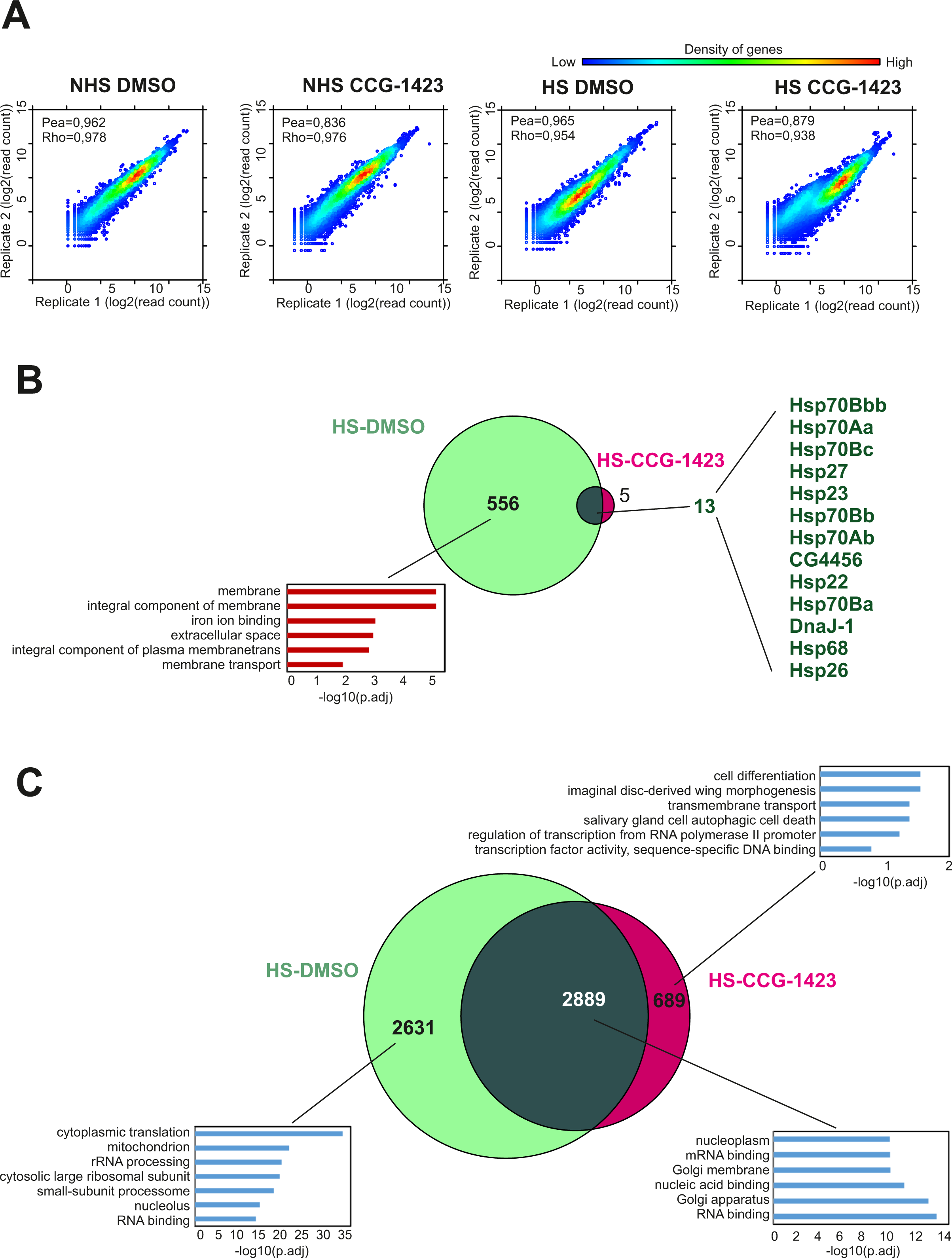
Correlation between PRO-seq replicates and differential gene expression upon HS and CCG-1423 treatments A. Correlation of spiked-in normalized read counts from two biological PRO-seq replicates at gene bodies of the Drosophila genome for NHS DMSO condition (pea=0,962, rhp=0,978), for NHS CCG-1423 condition (pea=0,836, rhp=0,976), for HS DMSO condition (pea=0,965, rhp=0,954), for HS CCG-1423 condition (pea=0,879, rhp=0,938). B. Venn diagram showing the overlap of genes induced upon HS in DMSO and CCG-1423 treated cells. Enriched gene ontology terms are indicated for the HS-induced genes and gene names for the common genes. C. Venn diagram showing the overlap of genes repressed upon HS in DMSO and CCG-1423 treated cells. Enriched gene ontology terms are indicated for the three classes of genes.

**Figure S7.**
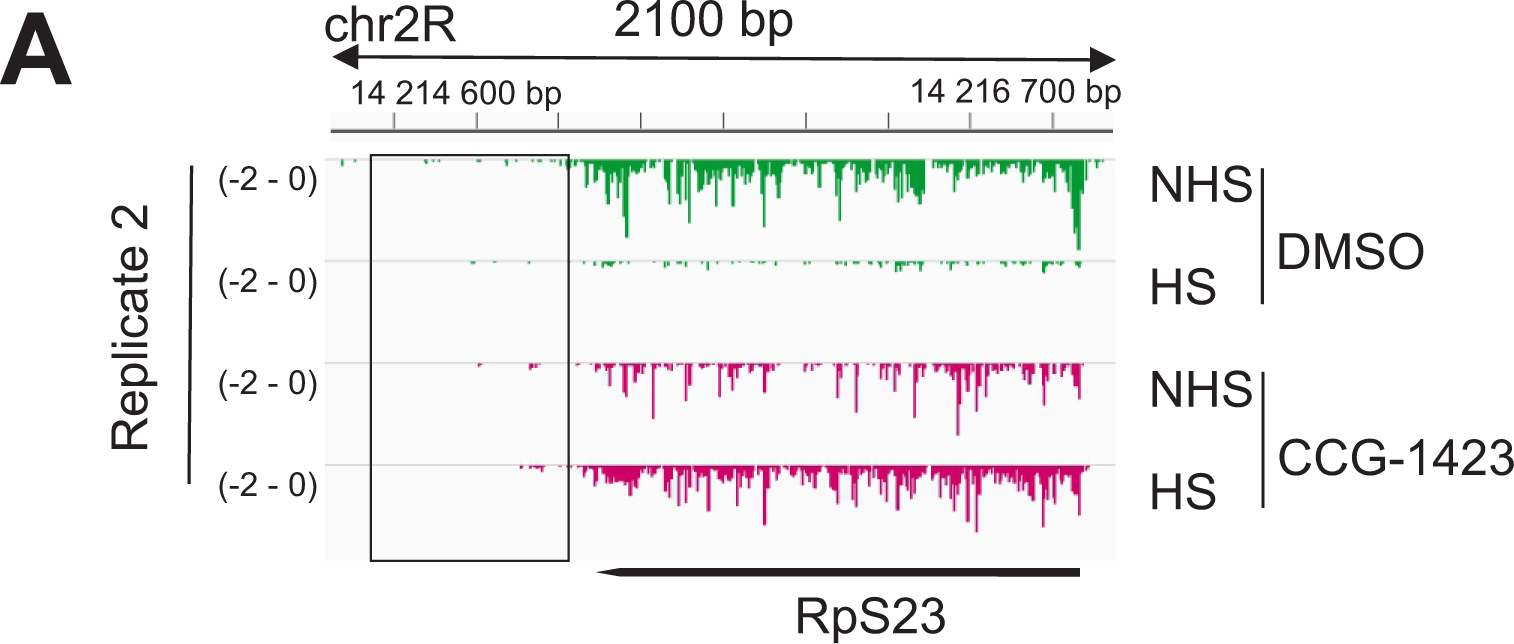
CCG-1423 reduces transcriptional repression upon heat shock **A.** Transcriptional profile of *Rps23* gene in S2R+ fly cells in non-heat shock (NHS) and heat shock (HS) conditions in DMSO or CCG-1423 treated cells (replicate2). Replicate 1 shown in Figure 7A.

**Figure S8.**
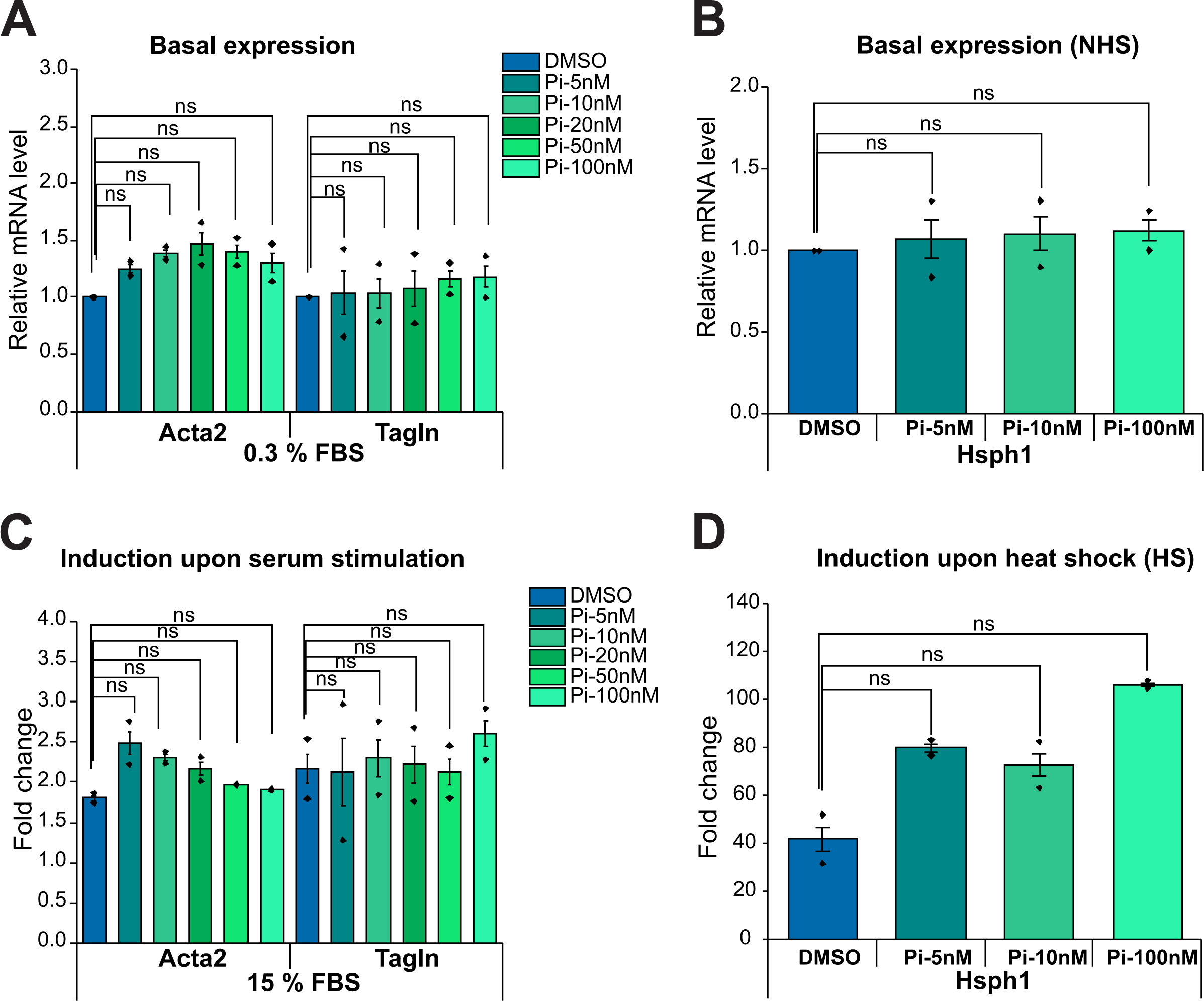
Pirin inhibitor CCT251236 does not influence heat shock or serum stimulated transcription A. Baseline expression of Acta2 and Tagln in NIH 3T3 cells upon treatment with pirin inhibitor CCT251236 for 10 mins. Bars represent fold change in Acta2 and Tagln mRNAs from two biological replicates with different concentrations of the inhibitor and untreated (DMSO) cells. Error bars are the standard error of mean (s.e.m.). Statistical significance measured with Student’s t-test (*; P< 0.01 with Bonferroni correction). P-values for Acta2: 5 nM CCT251236 vs DMSO is 0.0592, 10 nM CCT251236 vs DMSO is 0.0208, 20 nM CCT251236 vs DMSO is 0.1314, 50 nM CCT251236 vs DMSO is 0.0845, 100 nM CCT251236 vs DMSO is 0.2071. P-values for Tagln: 5 nM CCT251236 vs DMSO is 0.9292, 10 nM CCT251236 vs DMSO is 0.8936, 20 nM CCT251236 vs DMSO is 0.8287, 50 nM CCT251236 vs DMSO is 0.3563, 100 nM CCT251236 vs DMSO is 0.4319. B. Basal expression of Hsph1 in NIH3T3 cells treated with pirin inhibitor CCG251236 for 10 mins. Bars represent fold changes in the mRNA level of Hsph1 from two biological replicates with different concentrations of the inhibitor and untreated (DMSO) cells. Error bars are the standard error of mean (s.e.m.).Statistical significance measured with Student’s t-test (*; P< 0.017 with Bonferroni correction). P-values: 5 nM CCT251236 vs DMSO is 0.7973, 10 nM CCT251236 vs DMSO is 0.6669, 100 nM CCT251236 vs DMSO is 0.4131. C. Induction of Acta2 and Tagln upon inhibition of pirin in NIH 3T3 cells. Bars represent fold change in Acta2 and Tagln induction upon serum stimulation in cells treated with different concentrations of inhibitor and control (DMSO). Error bars are the standard error of mean (s.e.m.). Statistical significance measured with Student’s t-test (*; P< 0.01 with Bonferroni correction). For Acta2, P-values: 5 nM CCT251236 vs DMSO is 0.1325, 10 nM CCT251236 vs DMSO is 0.0325, 20 nM CCT251236 vs DMSO is 0.1521, 50 nM CCT251236 vs DMSO is 0.1084, 100 nM CCT251236 vs DMSO is 0.2493. For Tagln, P-values: 5 nM CCT251236 vs DMSO is 0.9644, 10 nM CCT251236 vs DMSO is 0.8436, 20 nM CCT251236 vs DMSO is 0.9376, 50 nM CCT251236 vs DMSO is 0.9390, 100 nM CCT251236 vs DMSO is 0.4715. D. Hsph1 induction in NIH 3T3 cells treated with pirin inhibitor CCT251236 under heat shock. Fold changes in Hsph1 mRNA induction in different concentrations of CCT251236 treated and untreated (DMSO) cells are calculated. Data is from two biological replicates and bars represent the mean value with individual data points as shown in E. Statistical significance measured with Student’s t-test (*; P<0.017 with Bonferroni correction). P-values: 5 nM CCT251236 vs DMSO is 0.0706, 10 nM CCT251236 vs DMSO is 0.1578, 100 nM CCT251236 vs DMSO is 0.0248.

**Figure S9.**
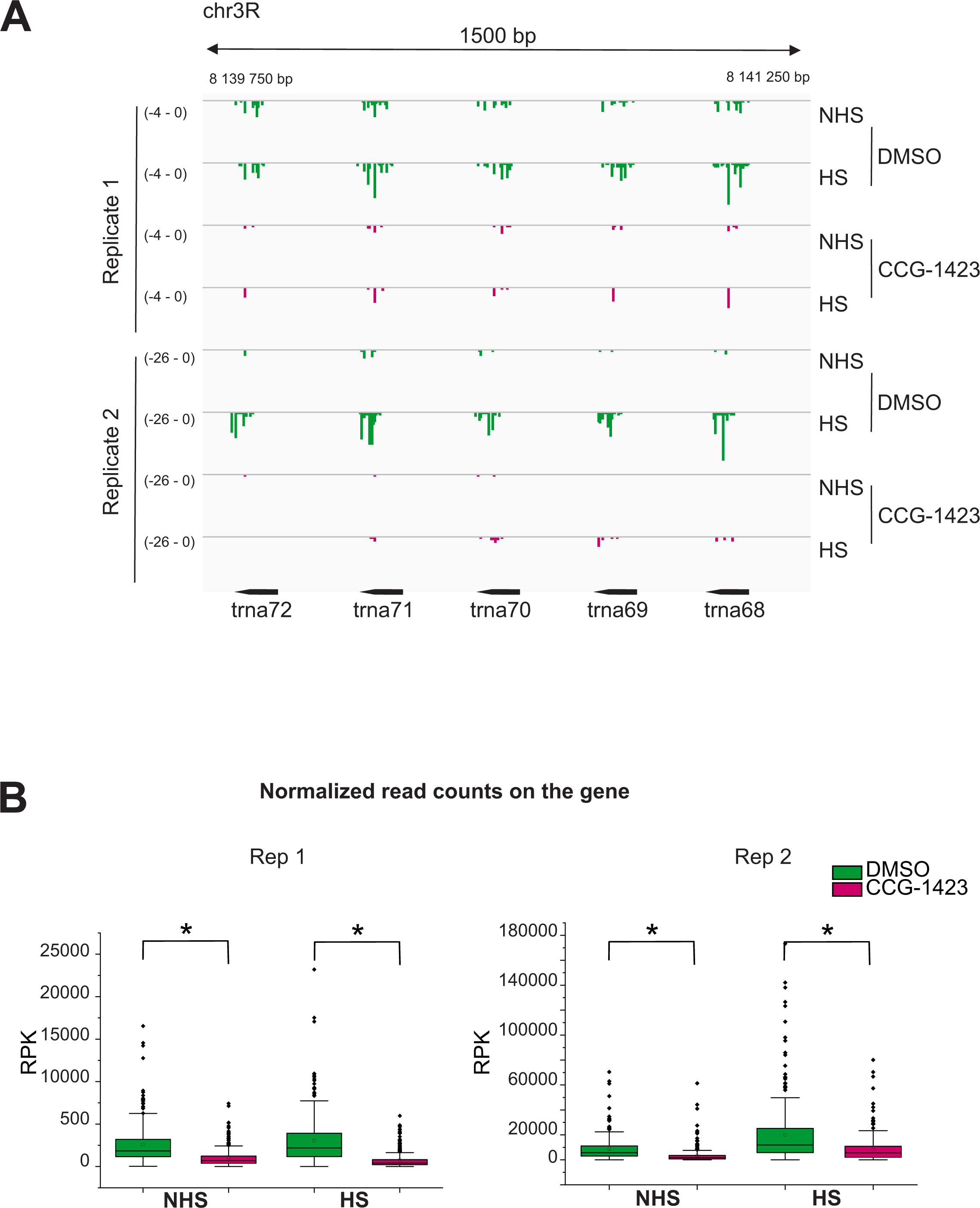
CCG-1423 reduced tRNA transcription **A.** Density of engaged Pol II (PRO-seq) at tRNA loci on the chr3R in S2R+ cells in non-heat shock (NHS) and heat shock (HS) conditions in DMSO or CCG-1423 treated cells.. **B.** Spike-in normalized read counts on 290 tRNA genes. Statistical significance was calculated by Mann-Whitney U test. (*; P< 0.05). P-values for NH DMSO vs CCG-1423: replicate 1 is 1E-00 and replicate 2 is 1E-00.

## References

Bell, J.L., A.J. Haak, S.M. Wade, P.D. Kirchhoff, R.R. Neubig, and S.D. Larsen. 2013. Optimization of novel nipecotic bis(amide) inhibitors of the Rho/MKL1/SRF transcriptional pathway as potential anti-metastasis agents. Bioorganic & medicinal chemistry letters. 23:3826–3832.

Berg, M.G., L.N. Singh, I. Younis, Q. Liu, A.M. Pinto, D. Kaida, Z. Zhang, S. Cho, S. Sherrill-Mix, L. Wan, and G. Dreyfuss. 2012. U1 snRNP determines mRNA length and regulates isoform expression. Cell. 150:53–64.

Booth, G.T., P.K. Parua, M. Sanso, R.P. Fisher, and J.T. Lis. 2018. Cdk9 regulates a promoter-proximal checkpoint to modulate RNA polymerase II elongation rate in fission yeast. Nature communications. 9:543.

Brandt, D.T., J. Xu, H. Steinbeisser, and R. Grosse. 2009. Regulation of myocardin-related transcriptional coactivators through cofactor interactions in differentiation and cancer. Cell Cycle. 8:2523–2527.

Cheeseman, M.D., N.E. Chessum, C.S. Rye, A.E. Pasqua, M.J. Tucker, B. Wilding, L.E. Evans, S. Lepri, M. Richards, S.Y. Sharp, S. Ali, M. Rowlands, L. O’Fee, A. Miah, A. Hayes, A.T. Henley, M. Powers, R. Te Poele, E. De Billy, L. Pellegrino, F. Raynaud, R. Burke, R.L. van Montfort, S.A. Eccles, P. Workman, and K. Jones. 2017. Discovery of a Chemical Probe Bisamide (CCT251236): An Orally Bioavailable Efficacious Pirin Ligand from a Heat Shock Transcription Factor 1 (HSF1) Phenotypic Screen. Journal of medicinal chemistry. 60:180–201.

Chu, T., E.J. Rice, G.T. Booth, H.H. Salamanca, Z. Wang, L.J. Core, S.L. Longo, R.J. Corona, L.S. Chin, J.T. Lis, H. Kwak, and C.G. Danko. 2018. Chromatin run-on and sequencing maps the transcriptional regulatory landscape of glioblastoma multiforme. Nat Genet. 50:1553–1564.

Cohen, R.S., and M. Meselson. 1985. Separate regulatory elements for the heat-inducible and ovarian expression of the Drosophila hsp26 gene. Cell. 43:737–746.

Cugusi, S., R. Mitter, G.P. Kelly, J. Walker, Z. Han, P. Pisano, M. Wierer, A. Stewart, and J.Q. Svejstrup. 2022. Heat shock induces premature transcript termination and reconfigures the human transcriptome. Mol Cell. 82:1573–1588 e1510.

Dennis, G., Jr., B.T. Sherman, D.A. Hosack, J. Yang, W. Gao, H.C. Lane, and R.A. Lempicki. 2003. DAVID: Database for Annotation, Visualization, and Integrated Discovery. Genome Biol. 4:P3.

Duarte, F.M., N.J. Fuda, D.B. Mahat, L.J. Core, M.J. Guertin, and J.T. Lis. 2016. Transcription factors GAF and HSF act at distinct regulatory steps to modulate stress-induced gene activation. Genes Dev. 30:1731–1746.

Evelyn, C.R., E.M. Lisabeth, S.M. Wade, A.J. Haak, C.N. Johnson, E.R. Lawlor, and R.R. Neubig. 2016. Small-Molecule Inhibition of Rho/MKL/SRF Transcription in Prostate Cancer Cells: Modulation of Cell Cycle, ER Stress, and Metastasis Gene Networks. Microarrays (Basel). 5.

Evelyn, C.R., S.M. Wade, Q. Wang, M. Wu, J.A. Iniguez-Lluhi, S.D. Merajver, and R.R. Neubig. 2007. CCG-1423: a small-molecule inhibitor of RhoA transcriptional signaling. Mol Cancer Ther. 6:2249–2260.

Galaxy, C. 2022. The Galaxy platform for accessible, reproducible and collaborative biomedical analyses: 2022 update. Nucleic Acids Res. 50:W345–W351.

Gau, D., P. Chawla, I. Eder, and P. Roy. 2022. Myocardin-related transcription factor’s interaction with serum-response factor is critical for outgrowth initiation, progression, and metastatic colonization of breast cancer cells. FASEB Bioadv. 4:509–523.

Geneste, O., J.W. Copeland, and R. Treisman. 2002. LIM kinase and Diaphanous cooperate to regulate serum response factor and actin dynamics. J Cell Biol. 157:831–838.

Guettler, S., M.K. Vartiainen, F. Miralles, B. Larijani, and R. Treisman. 2008. RPEL motifs link the serum response factor cofactor MAL but not myocardin to Rho signaling via actin binding. Mol Cell Biol. 28:732–742.

Haak, A.J., K.M. Appleton, E.M. Lisabeth, S.A. Misek, Y. Ji, S.M. Wade, J.L. Bell, C.E. Rockwell, M. Airik, M.A. Krook, S.D. Larsen, M. Verhaegen, E.R. Lawlor, and R.R. Neubig. 2017. Pharmacological Inhibition of Myocardin-related Transcription Factor Pathway Blocks Lung Metastases of RhoC-Overexpressing Melanoma. Mol Cancer Ther. 16:193–204.

Haeussler, M., K. Schonig, H. Eckert, A. Eschstruth, J. Mianne, J.B. Renaud, S. Schneider-Maunoury, A. Shkumatava, L. Teboul, J. Kent, J.S. Joly, and J.P. Concordet. 2016. Evaluation of off-target and on-target scoring algorithms and integration into the guide RNA selection tool CRISPOR. Genome Biol. 17:148.

Hayashi, K., B. Watanabe, Y. Nakagawa, S. Minami, and T. Morita. 2014. RPEL proteins are the molecular targets for CCG-1423, an inhibitor of Rho signaling. PLoS One. 9:e89016.

Henriques, T., D.A. Gilchrist, S. Nechaev, M. Bern, G.W. Muse, A. Burkholder, D.C. Fargo, and K. Adelman. 2013. Stable pausing by RNA polymerase II provides an opportunity to target and integrate regulatory signals. Mol Cell. 52:517–528.

Himanen, S.V., M.C. Puustinen, A.J. Da Silva, A. Vihervaara, and L. Sistonen. 2022. HSFs drive transcription of distinct genes and enhancers during oxidative stress and heat shock. Nucleic Acids Res. 50:6102–6115.

Johnson, L.A., E.S. Rodansky, A.J. Haak, S.D. Larsen, R.R. Neubig, and P.D. Higgins. 2014. Novel Rho/MRTF/SRF inhibitors block matrix-stiffness and TGF-beta-induced fibrogenesis in human colonic myofibroblasts. Inflamm Bowel Dis. 20:154–165.

Kallio, M.A., J.T. Tuimala, T. Hupponen, P. Klemela, M. Gentile, I. Scheinin, M. Koski, J. Kaki, and E.I. Korpelainen. 2011. Chipster: user-friendly analysis software for microarray and other high-throughput data. BMC Genomics. 12:507.

Kwak, H., N.J. Fuda, L.J. Core, and J.T. Lis. 2013. Precise maps of RNA polymerase reveal how promoters direct initiation and pausing. Science. 339:950–953.

Labun, K., T.G. Montague, M. Krause, Y.N. Torres Cleuren, H. Tjeldnes, and E. Valen. 2019. CHOPCHOP v3: expanding the CRISPR web toolbox beyond genome editing. Nucleic Acids Res. 47:W171–W174.

Laitem, C., J. Zaborowska, N.F. Isa, J. Kufs, M. Dienstbier, and S. Murphy. 2015. CDK9 inhibitors define elongation checkpoints at both ends of RNA polymerase II-transcribed genes. Nat Struct Mol Biol. 22:396–403.

Langmead, B., and S.L. Salzberg. 2012. Fast gapped-read alignment with Bowtie 2. Nat Methods. 9:357–359.

Lerdrup, M., J.V. Johansen, S. Agrawal-Singh, and K. Hansen. 2016. An interactive environment for agile analysis and visualization of ChIP-sequencing data. Nat Struct Mol Biol. 23:349–357.

Lisabeth, E.M., D. Kahl, I. Gopallawa, S.E. Haynes, S.A. Misek, P.L. Campbell, T.S. Dexheimer, D. Khanna, D.A. Fox, X. Jin, B.R. Martin, S.D. Larsen, and R.R. Neubig. 2019. Identification of Pirin as a Molecular Target of the CCG-1423/CCG-203971 Series of Antifibrotic and Antimetastatic Compounds. ACS Pharmacology & Translational Science. 2:92–100.

Love, M.I., W. Huber, and S. Anders. 2014. Moderated estimation of fold change and dispersion for RNA-seq data with DESeq2. Genome Biol. 15:550.

Lundquist, M.R., A.J. Storaska, T.C. Liu, S.D. Larsen, T. Evans, R.R. Neubig, and S.R. Jaffrey. 2014. Redox modification of nuclear actin by MICAL-2 regulates SRF signaling. Cell. 156:563–576.

Mahat, D.B., H. Kwak, G.T. Booth, I.H. Jonkers, C.G. Danko, R.K. Patel, C.T. Waters, K. Munson, L.J. Core, and J.T. Lis. 2016a. Base-pair-resolution genome-wide mapping of active RNA polymerases using precision nuclear run-on (PRO-seq). Nat Protoc. 11:1455–1476.

Mahat, D.B., and J.T. Lis. 2017. Use of conditioned media is critical for studies of regulation in response to rapid heat shock. Cell Stress Chaperones. 22:155–162.

Mahat, D.B., H.H. Salamanca, F.M. Duarte, C.G. Danko, and J.T. Lis. 2016b. Mammalian Heat Shock Response and Mechanisms Underlying Its Genome-wide Transcriptional Regulation. Mol Cell. 62:63–78.

Mali, P., L. Yang, K.M. Esvelt, J. Aach, M. Guell, J.E. DiCarlo, J.E. Norville, and G.M. Church. 2013. RNA-Guided Human Genome Engineering via Cas9. Science. 339:823–826.

Martin, M. 2011. Cutadapt removes adapter sequences from high-throughput sequencing reads. 2011. 17:3.

Medjkane, S., C. Perez-Sanchez, C. Gaggioli, E. Sahai, and R. Treisman. 2009. Myocardin-related transcription factors and SRF are required for cytoskeletal dynamics and experimental metastasis. Nat Cell Biol. 11:257–268.

Minami, T., K. Kuwahara, Y. Nakagawa, M. Takaoka, H. Kinoshita, K. Nakao, Y. Kuwabara, Y. Yamada, C. Yamada, J. Shibata, S. Usami, S. Yasuno, T. Nishikimi, K. Ueshima, M. Sata, H. Nakano, T. Seno, Y. Kawahito, K. Sobue, A. Kimura, R. Nagai, and K. Nakao. 2012. Reciprocal expression of MRTF-A and myocardin is crucial for pathological vascular remodelling in mice. EMBO J. 31:4428–4440.

Miralles, F., G. Posern, A.I. Zaromytidou, and R. Treisman. 2003. Actin dynamics control SRF activity by regulation of its coactivator MAL. Cell. 113:329–342.

Muniz, L., E. Nicolas, and D. Trouche. 2021. RNA polymerase II speed: a key player in controlling and adapting transcriptome composition. EMBO J. 40:e105740.

Posern, G., and R. Treisman. 2006. Actin’ together: serum response factor, its cofactors and the link to signal transduction. Trends Cell Biol. 16:588–596.

Rabenius, A., S. Chandrakumaran, L. Sistonen, and A. Vihervaara. 2022. Quantifying RNA synthesis at rate-limiting steps of transcription using nascent RNA-sequencing data. STAR Protoc. 3:101036.

Ramirez, F., D.P. Ryan, B. Gruning, V. Bhardwaj, F. Kilpert, A.S. Richter, S. Heyne, F. Dundar, and T. Manke. 2016. deepTools2: a next generation web server for deep-sequencing data analysis. Nucleic Acids Res. 44:W160–165.

Rawat, P., M. Boehning, B. Hummel, F. Aprile-Garcia, A.S. Pandit, N. Eisenhardt, A. Khavaran, E. Niskanen, S.M. Vos, J.J. Palvimo, A. Pichler, P. Cramer, and R. Sawarkar. 2021. Stress-induced nuclear condensation of NELF drives transcriptional downregulation. Mol Cell. 81:1013–1026 e1011.

Robinson, J.T., H. Thorvaldsdottir, W. Winckler, M. Guttman, E.S. Lander, G. Getz, and J.P. Mesirov. 2011. Integrative genomics viewer. Nat Biotechnol. 29:24–26.

Salvany, L., J. Muller, E. Guccione, and P. Rorth. 2014. The core and conserved role of MAL is homeostatic regulation of actin levels. Genes Dev. 28:1048–1053.

Sidorenko, E., M. Sokolova, A.P. Pennanen, S. Kyheroinen, G. Posern, R. Foisner, and M.K. Vartiainen. 2022. Lamina-associated polypeptide 2alpha is required for intranuclear MRTF-A activity. Scientific reports. 12:2306.

Sisson, T.H., I.O. Ajayi, N. Subbotina, A.E. Dodi, E.S. Rodansky, L.N. Chibucos, K.K. Kim, V.G. Keshamouni, E.S. White, Y. Zhou, P.D. Higgins, S.D. Larsen, R.R. Neubig, and J.C. Horowitz. 2015. Inhibition of myocardin-related transcription factor/serum response factor signaling decreases lung fibrosis and promotes mesenchymal cell apoptosis. Am J Pathol. 185:969–986.

Sokolova, M., H.M. Moore, B. Prajapati, J. Dopie, L. Merilainen, M. Honkanen, R.C. Matos, M. Poukkula, V. Hietakangas, and M.K. Vartiainen. 2018. Nuclear Actin Is Required for Transcription during Drosophila Oogenesis. iScience. 9:63–70.

Somogyi, K., and P. Rorth. 2004. Evidence for tension-based regulation of Drosophila MAL and SRF during invasive cell migration. Dev Cell. 7:85–93.

Tellier, M., J. Zaborowska, J. Neve, T. Nojima, S. Hester, M. Fournier, A. Furger, and S. Murphy. 2022. CDK9 and PP2A regulate RNA polymerase II transcription termination and coupled RNA maturation. EMBO Rep. 23:e54520.

Vartiainen, M.K., S. Guettler, B. Larijani, and R. Treisman. 2007. Nuclear actin regulates dynamic subcellular localization and activity of the SRF cofactor MAL. Science. 316:1749–1752.

Vihervaara, A., F.M. Duarte, and J.T. Lis. 2018. Molecular mechanisms driving transcriptional stress responses. Nat Rev Genet. 19:385–397.

Vihervaara, A., D.B. Mahat, M.J. Guertin, T. Chu, C.G. Danko, J.T. Lis, and L. Sistonen. 2017. Transcriptional response to stress is pre-wired by promoter and enhancer architecture. Nature communications. 8:255.

Vihervaara, A., D.B. Mahat, S.V. Himanen, M.A.H. Blom, J.T. Lis, and L. Sistonen. 2021. Stress-induced transcriptional memory accelerates promoter-proximal pause release and decelerates termination over mitotic divisions. Mol Cell. 81:1715–1731 e1716.

Vilborg, A., N. Sabath, Y. Wiesel, J. Nathans, F. Levy-Adam, T.A. Yario, J.A. Steitz, and R. Shalgi. 2017. Comparative analysis reveals genomic features of stress-induced transcriptional readthrough. Proc Natl Acad Sci U S A. 114:E8362–E8371.

Welch, W.J., and J.P. Suhan. 1985. Morphological study of the mammalian stress response: characterization of changes in cytoplasmic organelles, cytoskeleton, and nucleoli, and appearance of intranuclear actin filaments in rat fibroblasts after heat-shock treatment. J.Cell Biol. 101:1198–1211.

Wissink, E.M., A. Vihervaara, N.D. Tippens, and J.T. Lis. 2019. Nascent RNA analyses: tracking transcription and its regulation. Nat Rev Genet. 20:705–723.

Yu-Wai-Man, C., B. Spencer-Dene, R.M.H. Lee, K. Hutchings, E.M. Lisabeth, R. Treisman, M. Bailly, S.D. Larsen, R.R. Neubig, and P.T. Khaw. 2017. Local delivery of novel MRTF/SRF inhibitors prevents scar tissue formation in a preclinical model of fibrosis. Scientific reports. 7:518.

Zhang, Y., D. Song, Z. Peng, R. Wang, K. Li, H. Ren, X. Sun, N. Du, and S.C. Tang. 2022. LINC00891 regulated by miR-128-3p/GATA2 axis impedes lung cancer cell proliferation, invasion and EMT by inhibiting RhoA pathway. Acta Biochim Biophys Sin (Shanghai). 54:378–387.

Zhao, L., C. Li, W. Jiang, H. Luan, J. Zhao, J. Zhang, and Y. Xu. 2020. Serum response factor increases renal cell carcinoma migration and invasion through promoting epithelial-mesenchymal transition. Int J Urol. 27:808–816.

